# Data-driven analysis of fine-scale badger movement in the UK

**DOI:** 10.1101/2025.02.14.638245

**Authors:** Jessica Furber, Richard J. Delahay, Ruth Cox, Rosie Woodroffe, Maria O’Hagan, Naratip Santitissadeekorn, Stefan Klus, Giovanni Lo Iacono, Mark A. Chambers, David J.B. Lloyd

## Abstract

Understanding animal movements at different spatial scales presents a significant challenge as their patterns can vary widely from daily foraging behaviours to broader migration or territorial movements. This challenge is of general interest because it impacts the ability to manage wildlife populations effectively. In this study, we conduct diffusion analysis based on European badger (*Meles meles*) movement data obtained from three different regions in the UK (Gloucestershire, Cornwall, and Northern Ireland) and fit a generalised linear mixed-effects model to examine the relationship between variables. We also feature a novel application of *extended dynamic mode decomposition* (EDMD) to uncover patterns relating to badger social organisation. By applying our approach to these different populations, we were able to assess its performance across a range of badger densities. A key result was that in some areas, EDMD clusters matched observed group home ranges, whilst in others, discrepancies likely arose because of population management interventions, such as badger culling. The methods presented offer a promising approach for studying territoriality and the impacts of management strategies on animal movement behaviour.

**Author summary:** Wild animals move for many reasons, such as searching for food, finding mates, or maintaining territories, and these movements can be affected by changes in their environment. In this comparative study, we focused on European badgers, a social species whose movements are important for understanding behaviour and disease spread. Using GPS data from different locations around the UK, we explore how badger movement patterns vary both from day to day and over longer periods, revealing differences by sex, season, and region. The core contribution of this study lies in the novel application of extended dynamic mode decomposition (EDMD) alongside a more established generalised linear mixed-effects model (GLMM), together capturing movement dynamics across multiple timescales. These tools offer a new way to interpret animal behaviour including territoriality, and can be adapted to other species and ecological contexts. While dependent on tracking data, this approach lays the groundwork for future tools that may inform wildlife management and policy in conservation and disease control.

## Introduction

A fundamental challenge in the study of ecological processes, such as animal migration, swarming, and disease spread [1–3], is the identification and interpretation of movement patterns across different temporal scales. Short-term movement patterns, typically associated with daily activities like foraging, involve frequent changes in direction and speed. These fast temporal scale dynamics reflect the animal’s real-time interactions with its environment. In contrast, long-term or slow-scale patterns, such as migration or territory establishment, represent broader behavioural trends shaped over time. Understanding both scales is crucial, as it allows researchers to connect immediate behavioural responses with overarching ecological strategies, offering a more comprehensive view of animal movement and its drivers.

To model such patterns, researchers often employ stochastic differential equations (SDEs), which allow the simulation of animal positions in two-dimensional space. These models may incorporate spatial features, such as energy potentials representing gradients in resources or environmental conditions that attract or repel animals [4, 5], or may use individual-based stochastic simulations [6]. A key strength of the SDE framework is its natural separation of temporal scales: the *diffusion* term captures fast, random movements, while the *drift* term reflects slower, deterministic trends. For instance, high diffusion values might indicate erratic or exploratory movements during foraging, while the drift component may point to consistent directional movements due to resource availability or social behaviour. These terms also help reveal how movement patterns vary with factors such as season or sex (e.g., increased male ranging during mating periods). Additionally, certain areas where animals remain for prolonged periods, known as metastable states [7, 8], can be identified, offering insights into ecologically or behaviourally significant locations.

Our focal species for this investigation is the European badger (*Meles meles*), whose movement ecology and social organisation have been widely studied [9–13]. Badgers are behaviourally distinctive among mustelids, often forming relatively large, mixed-sex social groups and defending shared territories, though in some parts of their geographical range they are more solitary [9, 14]. Their ecological significance is further heightened by their role in the transmission of *Mycobacterium bovis*, the causative agent of bovine tuberculosis (bTB) [15–20]. Although cattle are the primary hosts, several wild mammals, including badgers, can contract and spread the disease [21], with evidence suggesting bidirectional transmission between badgers and cattle [22].

Despite ongoing uncertainties around the precise transmission routes, increasing attention has been paid to environmental contamination via badger foraging and defecation behaviour [18, 19, 23], including recent network analyses that underscore the importance of indirect contact at latrines [24]. Consequently, in both the United Kingdom and the Republic of Ireland, badgers have been subject to various management interventions, such as culling and vaccination [25, 26]. These interventions, especially culling, can disrupt social structures and potentially alter movement and disease dynamics [27–30].

Recent advances in movement ecology, including the use of GPS collars and high-resolution dead-reckoning data, have provided new insights into how badger behaviour responds to factors like cattle presence [19] and road construction [20]. While dead-reckoning allows for detailed analysis of fine-scale movements, it is typically limited to short durations due to data storage and battery constraints [31].

Hence, in the present study, we leverage long-term GPS tracking data from several field sites across the United Kingdom to characterise both short-term (diffusion) behavioural patterns and long-term (drift) movement trends in badgers. The sample of study populations represents a range of local badger densities and includes some locations where substantial temporal change in movement behaviour seems likely because of culling. The variations in conditions across our study populations provide an excellent opportunity to test the broad utility of our novel analytical approach. While our analysis provides ecological insights into badger behaviour, the broader value of the work lies in the generalisability of the method: our approach is adaptable to other species and landscapes, and has potential applications in understanding territoriality, space use, and behavioural response to environmental change.

## Methods and materials

In this section, we first give a brief introduction to the stochastic model, which the methods described are based on. Then, we progress to the mathematical methods, starting with estimation of the diffusion value from the data to gain further understanding of the dynamics of the everyday movements of badgers. This is followed by an introduction to the Koopman operator and extended dynamic mode decomposition, which make use of the trajectory data to estimate the number of metastable states present in the data. Then we explore the data collection, followed with the data cleaning and preparation for each method.

### The model

At time *t*, the position of a badger is given by **X**_*t*_ = (*x*_*t*_, *y*_*t*_)^⊤^ ∈ ℝ^2^ and can be modelled by a stochastic differential equation defined as

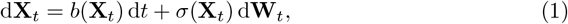

where *b* : ℝ^2^ → ℝ^2^ represents the *drift term, σ* : ℝ^2^ → ℝ^2×2^ is a real matrix that represents the *diffusion term*, and **W**_*t*_ is a two-dimensional Wiener process. The Wiener process (alternatively called Brownian motion) is widely used, typically to describe a random, but continuous, motion of atoms or particles, but here has been extended to model the movements of animals. We assume that badgers move isotropically in space, i.e. the diffusion is the same in all directions. This implies that we can write the diffusion matrix as a constant multiplied by the identity matrix, i.e., for some value *c* ∈ ℝ we can represent the diffusion matrix as

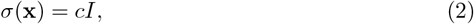

where *I* ∈ ℝ^2×2^ is the identity matrix.

### Estimating diffusion of badgers

The diffusion term in Eq. (1) can be estimated using the Kramers–Moyal formula, given by

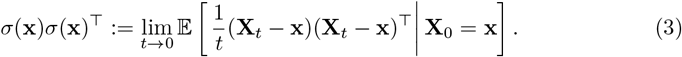

Because the data are collected as a time-series with discrete intervals, it is not possible to consider the limit as *t* → 0 in Eq. (3) (i.e., infinitely small time steps). However, an estimate can still be obtained using finite difference approximations. Since we assume that badgers move isotropically in space, we wish to calculate *c* in Eq. (2).

We estimate *c* as follows: given *m* measurements **x**_*i*_ = (*x*_*i*_, *y*_*i*_)^⊤^ at time *t*_*i*_, with *i* = 1, …, *m*, let Δ*t*_*i*_ = *t*_*i*+1_ − *t*_*i*_ denote the time difference between two successive GPS measurements, then

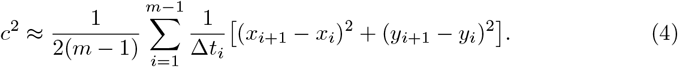

Numerical investigations show that good approximations to *c* and thus *σ* can be found despite ignoring the limit and when Δ*t*_*i*_ is not constant. Since *c* is always a positive number (as we take the square root), it is found that *c* is log-normally distributed, which is confirmed when running a Shapiro–Wilk test for normality [32].

We fitted a generalised linear mixed-effects model (GLMM) to examine the relationship between the diffusion value *c* and various predictors including capture year (i.e. GPS-collar monitoring year), month, and sex. The model also included a random intercept for animals-within-site to account for repeated measurements on the same individuals. The model used a normal distribution with a log link function and was fit using penalised maximum likelihood estimation (PL). Statistical tests were implemented in Matlab [33].

### Koopman operator and extended dynamic mode decomposition

We utilise the Koopman operator framework to analyse the dynamic behaviour of badger populations. We refer to the badger data as a dynamical system, which changes state over time according to a fixed rule, often described by differential equations such as Eq. (1). The eigenvalues and eigenfunctions of the Koopman operator provide valuable insights into the behaviour and stability of these systems. We are interested in applying the Koopman operator to badger movement data to investigate metastability, which refers to the tendency of the population to remain in a region for extended periods before transitioning to an alternative region.

We consider discrete trajectories from Eq. (1) with a state space ℝ^2^, representing the two-dimensional geographic location of a badger. The evolution of these trajectories is governed by an operator (also referred to as the flow map) **F** : ℝ^2^ → ℝ^2^ associated with the underlying dynamical system. In this context, the flow map describes how a badger’s position changes over time, essentially, it models how a badger moves from one location to another. More concretely, if **x**_0_ is the badger’s position at an initial time (e.g., the first GPS collar reading), then applying the flow map gives its estimated position after a fixed period of time *τ* :

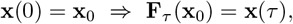

where *τ* represents a chosen time lag. This lag time can be interpreted as the interval between GPS observation, assuming those intervals are regular and consistent.

While the flow map **F** describes how a badger’s position evolves over time, the Koopman operator provides a complementary perspective by describing how measurable features of the system, such as speed, evolve. That is, whilst **F** acts directly on positions in the state space (i.e., **x** ∈ ℝ^2^), the Koopman operator 𝒦 acts on functions defined over the state space. We define the Koopman operator 𝒦_*τ*_ for data from a stochastic process as

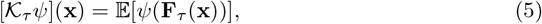

where **x** is an arbitrary point, 𝔼[·] denotes the expected value, and *ψ* ∈ *L*_∞_(ℝ^2^) is a measurable quantity of the system such as the speed of a badger. Under certain conditions, the operator can be extended from *L*_∞_ to *L*_2_ (i.e. so that it applies to a broader set of functions), allowing inner products to be used in Galerkin-like methods (see [34] for more details). Since the operator acts on functions, 𝒦_*τ*_ is infinite dimensional and linear even if **F** is finite dimensional and nonlinear.

Koopman analysis tries to find key measurement functions. The eigenfunctions *φ*(**x**) and the corresponding eigenvalues *λ* of the Koopman operator, defined by

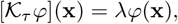

provide detailed insights into the underlying patterns and temporal evolution of the dynamical system. The eigenvalues *λ* ≈ 1 and the associated eigenfunctions are of particular interest in the badger movement system, as they provide crucial insights into the slow dynamics and the metastable sets. Small eigenvalues govern faster time scales, such as the daily dynamics of the badgers. The number of dominant eigenvalues corresponds to the number of metastable sets within the system. In some cases, it can be challenging to determine the number of metastable states due to no clear spectral gap, which is the barrier between the faster and slower dynamical modes. That is, given *N* eigenvalues *λ*_1_ *> λ*_2_ *>* … *> λ*_*N*_, a spectral gap is defined as the first *λ*_*i*_ such that

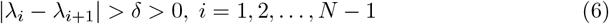

for some value of *δ >* 0. See S3 Appendix, where we plot the eigenvalues and their successive differences against *i* and also a different method to find the spectral gap to compare.

Extended dynamic mode decomposition (EDMD) is a data-driven method to estimate the Koopman operator and its eigenvalues, eigenfunctions, and modes. The method requires a data set of snapshot pairs, i.e.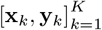, where each pair would consist of observations of the dynamical system such that **y**_*k*_ = **F**(**x**_*k*_). In other words, given time series data **x**_*k*_, then **y**_*k*_ = **F**(**x**_*k*_) is the next observation in the time series. Each observation is a GPS-derived location of a badger, numbers of which vary among sites (*n*), from which a finite number of data points are collected *m*_*i*_ (for *i* = 1, …, *n*), and the value of *m* varies from animal to animal. The coordinate data for the *i*th badger is represented as a vector, 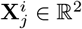 at time *t*_*j*_, where *j* = 1, …, *m*_*i*_. Ideally, the time intervals between observations in the time series are equal, as EDMD assumes evenly spaced data for accurate analysis. When the time intervals are irregular, interpolation is necessary to construct a uniformly spaced time series before applying the method. We organised the GPS data in the matrices 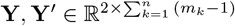,

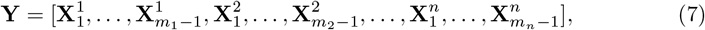

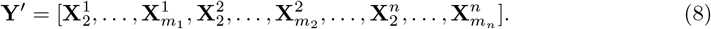

In simpler terms, the **Y** matrix is made up of the first pair of coordinates to the penultimate coordinates for each badger, and the **Y**′ matrix starts from the second entry of coordinates and includes all entries up to the last.

The method also requires a dictionary of observables *ψ*, 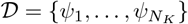, where *N*_*K*_ (the number of basis functions) is always finite-dimensional in practice because working with infinite-dimensional dictionaries is not feasible [34]. The function space itself, however, remains infinite-dimensional. Consequently, the operator learned from the data is an approximation, specifically a Galerkin projection. Here, indicator functions are used as the set of *basis functions*—a mathematical term referring to foundational building blocks for approximating complex functions—in EDMD, which is equivalent to using Ulam’s method to estimate a Markov state model [35]. Other basis functions such as monomials up to a certain order could be used. However, the resulting matrices might then become numerically ill-conditioned. The optimal choice of dictionary elements depends strongly on the dynamical system and remains an open question. When defining the domain for the EDMD indicator functions, we considered a box discretisation (i.e. we superimposed a grid over the area of coordinates). For consistency, the domain was divided into a fixed number of grid cells, requiring the number of boxes along each axis to be an integer. This resulted in box dimensions that were approximately 100 m by 100 m, with minor variation across years depending on the extent of the domain.

Let 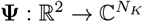 be the vector-valued function

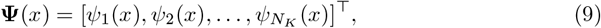

for an arbitrary value of *x*. Then, we define the transformed data matrices 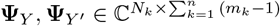 as

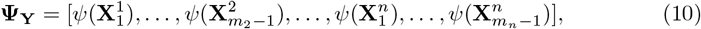

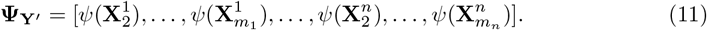

The minimisation problem for the transformed data matrices becomes,

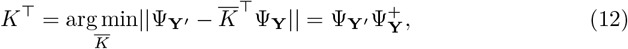

where ^+^ denotes the Moore–Penrose pseudoinverse of a matrix. The matrix 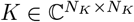 is a representation of the Koopman operator 𝒦_*τ*_ projected onto the subspace spanned by the chosen basis functions, i.e. a Galerkin approximation.

Finally, let *ξ*_*j*_ be the *j*th eigenvector of *K* with the eigenvalue *λ*_*j*_, then the EDMD approximation of an eigenfunction of 𝒦_*τ*_ is

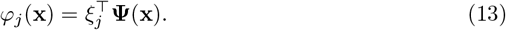

Once the Koopman eigenvalues and eigenfunctions have been computed, the number of dominant eigenvalues is chosen based on the spectral gap defined in Eq. (6) and the corresponding eigenfunctions are clustered using *k*-means clustering [36]. For example, if there are eight dominant eigenvalues, the corresponding first eight eigenfunctions form clusters that define metastable states, representing the spatial areas that badgers rarely leave.

### Data collection

We accessed three sets of data from studies where collars carrying GPS devices had been fitted to badgers. One dataset derived from a study of badgers resident in Woodchester Park, Gloucestershire, England [13, 23], another related to four different sites in Cornwall, England [19], and the third was from a population in County Down, Northern Ireland [20].

The Woodchester Park Study was established in 1976 to investigate TB in a wild, naturally infected badger population [37]. The study includes analysis of capture-mark-recapture data, which involves trapping and re-trapping individually marked (tattooed) badgers to infer group membership [13], and the analysis of GPS data. The study site itself covers *c*. 7 km^2^ of predominantly permanent pasture and woodland [12]. When designing the data collection schedule at Woodchester, a 35-minute interval was chosen as a balance between collecting sufficient GPS fixes and preserving collar battery life. The collar was programmed to collect fixes from 20:00 to 04:00 daily. Due to the collar’s internal scheduling based on a midnight-to-midnight cycle, the last fix of one day is taken at 23:30, and the first fix of the next day is taken at 00:00. While this creates a 30-minute interval between these two fixes, it is a consequence of the scheduling boundary at midnight, not an intentional deviation from the 35-minute interval.

The four sites in Cornwall each included at least two beef farms and two dairy farms and were chosen to achieve representation of varying environmental conditions [19]. Sites C2 and C4 in North Cornwall, and F2 on the South Coast of West Cornwall are dominated by pastoral landscapes with wooded valleys, and site F1 on the North coast of West Cornwall is bounded by granite cliffs and moorland. The GPS units deployed on badgers recorded position fixes every 20 minutes between 18:00 and 06:00 while the badger was active.

The Northern Ireland study area in County Down, near the town of Banbridge, covers *c*. 100 km^2^ of primarily pasture and arable land [38]. The resident badger population was the subject of a Test and Vaccinate or Remove (TVR) project whereby captured animals were tested for TB using the Dual-Path Platform VetTB test (DPP) and test-positives were euthanised, whilst test-negative badgers were vaccinated against TB with Bacille Calmette Guerin, fitted with GPS collars and released [38–40]. The GPS units recorded position fixes once per hour between 21:00 and 04:00.

The various study areas contained different numbers of badger social groups and individuals fitted with GPS devices over a range of time periods (Table 1). The number of social groups was estimated using field knowledge specific to each site: in Woodchester, bait-marking data were used [41]; in Northern Ireland and Cornwall, capture-recapture and GPS collaring data were applied (Table 1 shows the total number of social groups monitored throughout the total GPS-collar monitoring period). GPS data in the form of locational fixes were not captured at a uniform rate within or among years of study (Fig 1), although most fixes were collected in the latter half of the year. Most collars were deployed in the summer/autumn (the peak season for badger trapping [42]), which means that the battery life waned during the first half of the following year, which overlaps with the closed season when trapping is suspended to prevent disturbance of reproductive females and their offspring [43]. Battery longevity is dependent on the frequency with which fixes are recorded. The temporal distribution of captured badgers (split by sex) for each site is shown in S3 Fig, highlighting variability in the number of individuals monitored (i.e., with active GPS collars) per month. At their respective peaks, Northern Ireland monitored twice as many animals as Woodchester, and up to six times more than the Cornwall sites.

**Table 1.** Summary of site and badger monitoring across all study sites and years used in this study.

| Site | Woodchester | C2 | C4 | F1 | F2 | Northern Ireland |
| --- | --- | --- | --- | --- | --- | --- |
| Approximate Size | *c.*7 km<sup>2</sup> | *c.*8 km<sup>2</sup> | *c.* 6 km<sup>2</sup> | *c.*8 km<sup>2</sup> | *c.*6 km<sup>2</sup> | *c.*100 km<sup>2</sup> |
| Years Monitored | 2018–2022 | 2013–2015 | 2014–2017 | 2013–2017 | 2013–2015 | 2014–2017 |
| Social Groups | 18 | 6 | 5 | 7 | 10 | 45 |
| *Badgers GPS-Collared* |  |  |  |  |  |  |
| Total | 66 | 12 | 13 | 22 | 20 | 105 |
| Female | 34 | 3 | 9 | 13 | 12 | 41 |
| Male | 32 | 9 | 4 | 9 | 8 | 64 |

**Fig 1.**
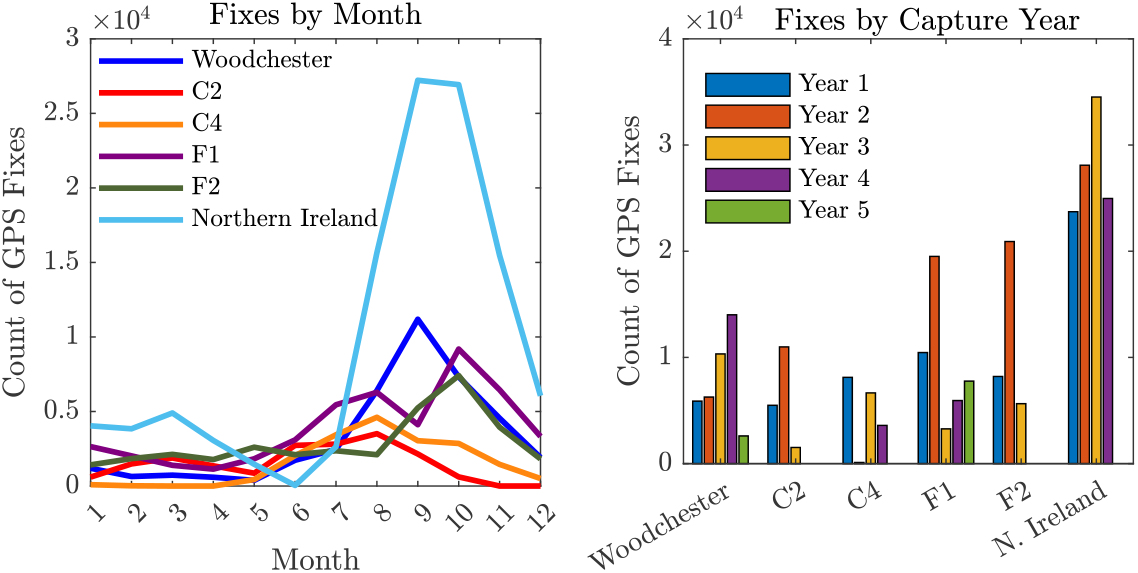
Summary of distribution of GPS fixes. Number of fixes by month (left) and by capture year for respective sites (right).

To account for the impact of culling and other interventions across the different study sites, it is important to consider their varying management histories. At Woodchester, badger culling began on the farmland surrounding the core of the study area in 2018 (Woodchester Park belongs to the National Trust which has a no-cull policy), meaning one might expect change to the social structure in all years from 2018 onwards due to population reductions. In Cornwall, there was no active culling during the study period for sites F1, F2 and C2. Although site F1 was in a former treatment area of the Randomised Badger Culling Trial (RBCT) [44], potentially influencing badger dynamics through residual effects from past culling. The Cornwall site C4, however, was culled from 2016 onwards [18]. In Northern Ireland, the selective removal of TB-positive badgers after the first year of the study [38] might have disrupted social cohesion, prompting increased ranging behaviour or social group reconfigurations in response to changing population density. These variations in culling and TB management strategies must be considered when interpreting EDMD results and comparing them with field observations of badger social groups, as they likely played a role in the movement patterns observed.

### Data cleaning and preparation

To avoid location errors, and to be consistent with the pre-cleaned data obtained from the Cornwall sites [19], we excluded all GPS-collar locations associated with fewer than four satellites, or with *dilution of precision* greater than four (as detailed in Supporting Information [25]) from the Woodchester and Northern Ireland data sets (where the parameters allowed). Moreover, improbable GPS locations (e.g., those far outside the study areas, or in water bodies), were excluded from all data sets. We chose to remove data derived from one Woodchester Park badger and five individuals from the Northern Ireland study because they were clearly isolated from the remaining data (see S1 Fig). In line with previous studies where badgers are recorded to have been moving at speeds of up to 26.2 m min^−1^ [11], we excluded any data points where badgers were found to be moving faster. Additionally, any single data points where the recorded movement within a time interval appeared biologically unrealistic for badgers were removed, such as travelling a distance of greater than 1500 m in two consecutive time steps (e.g. 70 minutes). As a result of these actions, the Woodchester dataset was reduced by 19.0%, the Northern Ireland data by 13.3%, and the data from the Cornwall sites was reduced by 1.1%. See S1 Appendix for the breakdown of reasoning for data removal.

The coordinates of the cleaned data were converted from the longitude/latitude (Woodchester and Northern Ireland) or British National Grid (Cornwall) into UTM (Universal Transverse Mercator) coordinates, so the distances between coordinates could be measured in meters. To preserve confidentiality of the location of sites (and to be consistent), the coordinates were shifted to a (0,0) base. For the estimation of the diffusion metric for badgers, the data can be used as presented. However, to account for the large distances between the last location of one night-time monitoring period and the first location of the next, during which time badgers were likely inactive in their underground setts, these intervals were excluded from the calculation. Furthermore, since the limit in Eq. (3) cannot be taken, once the individual diffusion metric had been estimated, it was decided to exclude any individual’s measurements derived from ten or fewer data points, as an insufficient number of points hinders the accurate estimation of the diffusion metric.

Continuous and evenly spaced time series data are essential for EDMD, as the method assumes uniform time intervals between observations. However, GPS data for badgers often features irregular intervals, necessitating interpolation to create consistent time steps. These inconsistencies can arise due to undetected signals (e.g., when badgers are underground or beneath dense vegetation), timing issues with GPS devices (e.g., a 70-second window to connect with satellites), or large gaps caused by GPS collars losing power, which can result in days or months without data until the collar is replaced. To manage these gaps, new ID numbers were assigned to separate series of GPS fixes, ensuring continuity in the data. For instance, at Woodchester, GPS data were available from 66 unique badgers, but this was increased to 108 collar monitoring periods after accounting for interruptions. Recorded times (e.g., 20:01) were adjusted to the nearest pre-programmed time point (e.g., 20:00) to standardise the timing of locations. Notably, the time discrepancy reflects the duration between the start of GPS location calculation and the time of recording, which can vary across sites. At Woodchester Park, likely influenced by its high proportion of woodland, 13.6% of badgers did not have their positions recorded at the designated time, with most locations logged within one minute of the intended capture time, but some taking up to three minutes. In contrast, only 5% of the badgers at the Cornwall sites failed to register their location within one minute. Most badgers in the Northern Ireland study area registered a location fix within sixty seconds of the appointed hourly time. Those that did not (see [45]) were excluded (2.5%) to facilitate interpolation. Although badgers are known to travel along complex paths (see [31]), linear interpolation was used to create evenly spaced intervals, closely matching the original time steps. This approach minimises the impact of interpolation on the data matrices used in EDMD.

For example, when locations were recorded every 35 minutes, such as in Woodchester, and a factor of 35 minutes is not chosen, we observe cutting corners and missing the actual recorded data point, resulting in calculating a shorter distance travelled by the badger. This underestimation is expected in any sampling of locations from continuous time, as linear interpolation inherently misses finer-scale movements and underestimates the true distance travelled. We found choosing a two-minute interpolation interval produced only small differences between consecutive positions, which were too subtle for EDMD to reliably detect underlying movement dynamics. At the same time, this high-frequency interpolation introduced artificial detail by generating multiple inferred locations between actual observations, thereby making more assumptions about the badger’s movement than were supported by the data. Using a 35-minute interval caused EDMD to fail in identifying non-trivial dynamics because the large time gap between points missed finer-scale transitions, oversimplified the system’s behaviour, and reduced the temporal resolution needed to capture complex, non-linear dynamics. For the Woodchester data, we found the optimal interval to be 17.5 minutes because it only assumes one location of the badger between observations, and EDMD was able to identify meaningful dynamics. For the Cornwall datasets, which comprised GPS fixes at 20-minute intervals, this was also found to be optimal. In contrast, the Northern Ireland dataset contained fixes recorded at hourly intervals, necessitating interpolation to create data points every 20 minutes for consistency with the Cornwall datasets. It is important to note that this approach means approximately two-thirds of the Northern Ireland dataset consists of interpolated values, which should be considered when interpreting the results.

## Results

### Badger diffusion

The results of the GLMM analysis investigating badger diffusion patterns are presented in Tables 2 and 3 (see S2 Appendix for univariate analyses and S2 Fig for the data inputted into the model). The model included two fixed predictors: month and sex. Their respective estimates, standard errors, and confidence intervals are reported in Table 2. The model also allows for random slopes to account for variation in capture years across sites.

**Table 2.** Fixed effect estimates from the final generalised linear mixed-effects model (GLMM) with a normal distribution and log link.

| Predictor | Estimate | Standard Error | t-Statistic | 95% Confidence Interval | p-Value |
| --- | --- | --- | --- | --- | --- |
| Intercept | 3.397 | 0.041 | 82.80 | [3.32, 3.48] | < 0.001 |
| Month (Ref: January) |  |  |  |  |  |
| February | 0.106 | 0.023 | 4.54 | [0.060, 0.15] | <0.001 |
| March | 0.050 | 0.026 | 1.89 | [-0.0018, 0.10] | 0.056 |
| April | 0.036 | 0.030 | 1.20 | [-0.023, 0.094] | 0.23 |
| May | 0.006 | 0.033 | 0.188 | [-0.059, 0.071] | 0.85 |
| June | -0.106 | 0.038 | -2.75 | [-0.18,-0.030] | 0.0060 |
| July | -0.069 | 0.027 | -2.56 | [-0.12, -0.016] | 0.011 |
| August | -0.116 | 0.021 | -5.39 | [-0.16, -0.074] | <0.001 |
| September | -0.236 | 0.021 | -11.09 | [-0.28,-0.19] | <0.001 |
| October | -0.370 | 0.023 | -16.34 | [-0.41, -0.33] | <0.001 |
| November | -0.442 | 0.024 | -18.36 | [-0.49, -0.40] | <0.001 |
| December | -0.290 | 0.024 | -11.87 | [-0.34, -0.24] | <0.001 |
| Sex (Ref: Male) |  |  |  |  |  |
| Female | -0.152 | 0.034 | -4.42 | [-0.22, -0.084 ] | <0.001 |

**Table 3.**
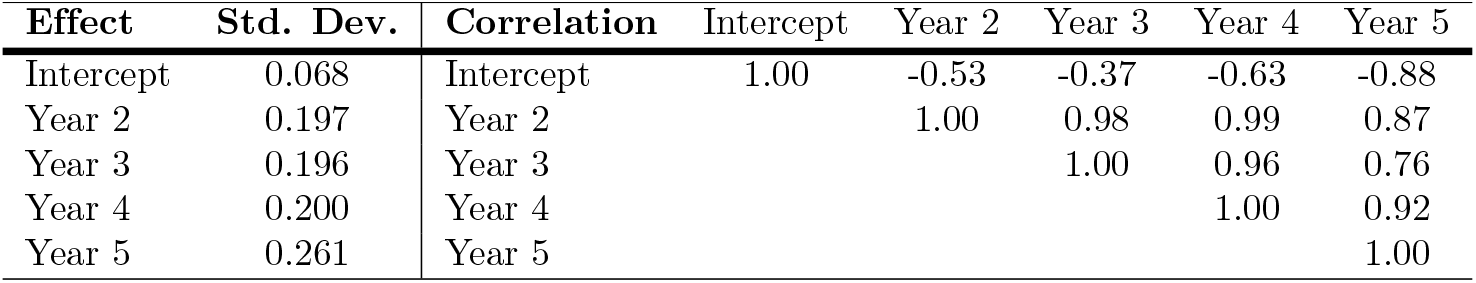
Random effects for Capture Year at the Site level. Standard deviations and correlations.

| Effect | Std. Dev. | Correlation | Intercept | Year 2 | Year 3 | Year 4 | Year 5 |
| --- | --- | --- | --- | --- | --- | --- | --- |
| Intercept | 0.068 | Intercept | 1.00 | -0.53 | -0.37 | -0.63 | -0.88 |
| Year 2 | 0.197 | Year 2 |  | 1.00 | 0.98 | 0.99 | 0.87 |
| Year 3 | 0.196 | Year 3 |  |  | 1.00 | 0.96 | 0.76 |
| Year 4 | 0.200 | Year 4 |  |  |  | 1.00 | 0.92 |
| Year 5 | 0.261 | Year 5 |  |  |  |  | 1.00 |

Alternative model structures were explored, including configurations with both temporal variables (month and capture year) modelled as random slopes. While these models maintained a good fit (R^2^ *>* 0.7), their performance declined relative to the final model, as indicated by higher AIC values and less interpretable variance structures. Model comparison statistics, including AIC and R^2^, are provided in S1 Table. Two models appeared to perform best: the model presented in Tables 2 and 3, which includes random slopes for capture year only, and the model without capture year (the last model in S1 Table). While the first model outperforms the second model in terms of AIC (80.819 vs. 152.82), the second model outperforms the first model in terms of BIC (238.66 vs. 231.74). Hence, the fixed effects estimates for the second model have also been reported, in S2 Table, where it can be observed that the estimates are similar to those observed in Table 2. The decision to choose the first model was supported by the fact that capture years differed across sites due to varying management regimes, introducing meaningful heterogeneity best captured with random slopes on year. Diagnostic plots for residuals of the final model are shown in S4 Fig.

Monthly variation in diffusion rate was evident, with reduced movement observed from June to December relative to January. Notably, movement increased in February, with diffusion rates approximately 11% higher than in January (exp(0.106) = 1.112, *p <* .001), suggesting an early seasonal shift in activity. Sex was also a significant predictor of diffusion rate. Female badgers exhibited a reduction in movement compared to males, with an estimated 14% lower rate (exp(−0.152) = 0.859, *p <* .001).

Random effects associated with capture year at the site level (Table 3) indicated modest variation in site-specific intercepts (SD = 0.068), but substantial variation in slopes across years (SD range = 0.197–0.261). This suggests that temporal trends differ across sites, some may exhibit increasing diffusion rates over time, while others may remain stable or decline. Additionally, we observe a strong correlation between capture years.

The random intercept variance for animals within sites was 0.229, reflecting individual-level heterogeneity in movement. The residual standard deviation 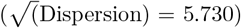 indicates that, despite the model’s complexity, unexplained variability in diffusion rates remains, likely attributable to unmeasured environmental or behavioural factors.

### Extended dynamic mode decomposition

We employed EDMD to analyse the drift term in the stochastic differential equation describing badger movement behaviour. By extracting dominant modes and eigenvalues from observed trajectories, EDMD effectively approximated the drift component, which represents the average or expected rate of change in the system’s state.

Fig. 2 displays the calculated metastable clusters for Woodchester across the full dataset and the individual years. Each panel shows badger location, with individual location fixes represented as circles, colour-coded according to their assigned metastable cluster. The purple lines represent the convex hulls of these metastable clusters, outlining the spatial extent of each cluster (with the area estimated next to it). The orange lines represent the convex hulls of social group home ranges derived directly from the raw movement data. A convex hull is the smallest convex polygon that contains all the points in the given set (calculated here using the *convhull* function in MATLAB). In this study, convex hulls provide a simple and interpretable way to define the spatial extent of badger activity within each cluster or social group. For further examples from the other sites, see S5 Fig.

**Fig 2.**
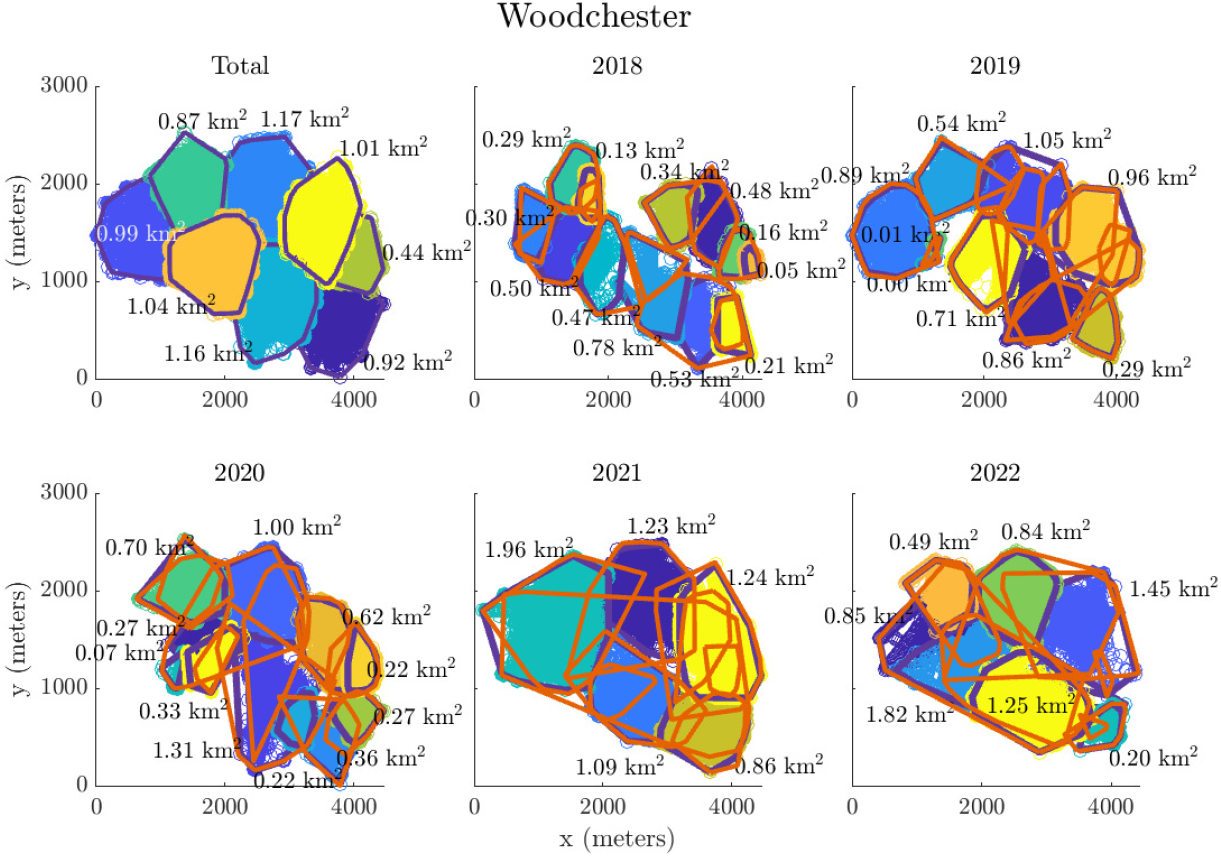
The calculated metastable clusters with areas of clusters for Woodchester. The purple lines are the convex hull of the metastable clusters, the orange lines are the convex hull of the social group movement (home ranges) based on the data.

Table 4 summarises the number of social groups identified at each site alongside the groupings calculated using EDMD. It also presents the mean home range area of the social groups and the mean area of the metastable clusters, both estimated from the area of their respective convex hulls. The amalgamated multi-year clustering provides a broad overview of the long-term trends in movement behaviour, where it can be observed that EDMD predicted fewer clusters than those observed via capture–mark–recapture. This pattern was most evident in Northern Ireland, where EDMD predicted 15 metastable clusters where 45 social groups had been identified by field observations, capture/recapture and GPS tracking.

**Table 4.**
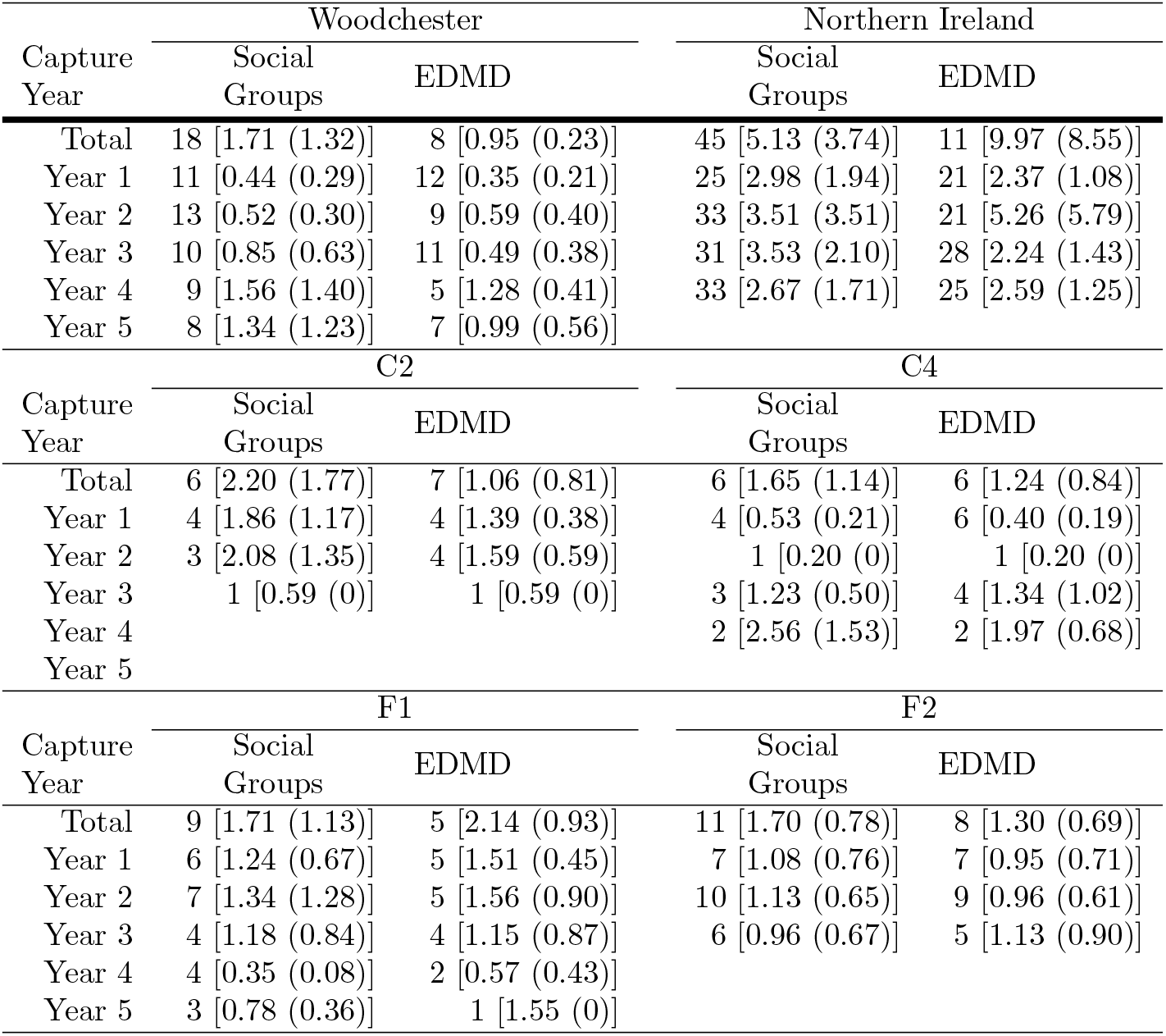
Summary of badger social groups at each site and capture year. For each site, the number of observed social groups (left column) is shown alongside the number of clusters identified using extended dynamic mode decomposition (EDMD; right column). Bracketed values represent the mean home range area or cluster area in km^2^ (standard deviation).

Although some clusters appeared to be stable for periods of time, the number of metastable states can be seen to vary per year. For example, in years two to five at Woodchester (Fig 2), a cluster persisted in the central north of the study area (covering 1.05 km^2^, 1.00 km^2^, 1.23 km^2^, and 0.84 km^2^, respectively). Yet, during those same years, the estimated number of clusters fluctuated between one and two in the northeast. Similarly, EDMD identified four metastable clusters in the west of the study area throughout the capture period, but this reduced to one large cluster in the fourth year. Similar observations can be made in other sites, for example, in the Cornwall site C4 (S5 Fig, C), both the first and second capture year present similar clustering. When there is only one social group present in the data for the year (C2 and C4 during capture year three and two, respectively) then EDMD rightly picks up only one metastable state.

## Discussion

Badgers are social animals that often live in groups occupying mutually defended territories across much of their geographic range [9], although extra-territorial movements are not uncommon [13]. Traditional ecological methods, such as bait-marking and capture-mark-recapture studies, have been invaluable for identifying group territories and changes in social structure [41, 46], while radio-tracking has provided further insights into individual movement patterns despite its labour-intensive nature and limited resolution [47]. The advent of GPS technology has since revolutionised the study of badger movements, offering finer-scale tracking that generates rich datasets suited to novel mathematical analyses.

Building on these advancements, our study applied a combination of diffusion analysis and EDMD to investigate the intricate movement patterns and social behaviours of badgers. This approach allowed us to characterise movement patterns across fast (diffusion) and slow (drift) spatiotemporal scales, while identifying effective metastable group home ranges over time. The analysis spanned GPS data from badger populations in three distinct regions of the United Kingdom, representing a range of local badger densities. Importantly, the study has included populations exposed to varying degrees of culling, providing a unique opportunity to examine the impact of external interventions on movement behaviour. The diversity in conditions across these populations underscores the utility of our novel analytical approach and highlights its potential to yield new insights into badger spatial organisation and movement dynamics.

We first examined the fast temporal dynamics from badger GPS data to gain an understanding of regular badger movements. Diffusion, a metric that captures both spatial extent (distance) and frequency of movement, was used to quantify these dynamics. Our results indicate that males exhibit a higher diffusion value than females, suggesting they have greater spatial mobility. This could mean they are covering larger distances, visiting more locations, or roaming across a broader area more frequently. In contrast, the movements of females appear more localised. Research on the regular, individual movements of male and female badgers has been limited, as the use of GPS tracking technology has only become widespread in the past decade. Prior to the introduction of GPS tracking, many studies focused on movement data collected between capture events, which provided only intermittent snapshots of badger activity [13]. These studies typically infer movement patterns based on locations obtained at specific intervals, which can miss finer details of regular movement dynamics.

From our results, monthly variations in badger movement are evident in the estimated diffusion value, especially during the latter half of the year (Table 2). An increase in movement during certain periods was also reported by Woodroffe et al. [48], who (using the same dataset from Cornwall sites), found clear peaks in the frequency of territorial trespassing in February and September. This trend is most apparent in Woodchester (S2 Fig, A), where increased movement during this time may reflect heightened activity related to reproductive strategies [49]. This observation is consistent with broader trends in wildlife movement patterns, where mating strategies and the need for reproduction can drive differences in spatial behaviours between the sexes. Furthermore, when compared to January (the reference month in the GLMM), the model suggests that movement patterns in the latter half of the year differ significantly, indicating seasonal shifts in behaviour. This reinforces the idea already seen (see [48]) that environmental and reproductive factors influence movement dynamics across different times of the year.

Moreover, diffusion rates differed between the years of capture, suggesting temporal variability in movement patterns. While the overall variance in site-level intercepts was modest, the inclusion of random slopes for capture year revealed substantial differences (Table 3; SD range: 0.196–0.261). While the model does not directly specify whether diffusion increased or decreased over time at individual sites, the variability suggests heterogeneous temporal trends, with some sites potentially showing increases, others showing stable or declining rates. These variations are likely influenced by site-specific factors such as habitat changes, management interventions, or local population dynamics, emphasizing the importance of accounting for temporal structure in the analysis. This finding aligns with recent research emphasising the significance of spatiotemporal context in understanding animal movement dynamics, where land-use change can significantly alter wildlife movement patterns and behaviour [50]. Though further analysis would be required to identify specific drivers for the different results.

The univariate analysis revealed a significant difference between the diffusion values across all sites, where it was highlighted through pairwise comparisons that the diffusion values at Woodchester and the Cornwall sites significantly differed from those from Northern Ireland. Existing research indicates that badger movement can vary with population density [51]. For instance, comparison of two populations identified higher movement frequency at lower-density and a more fluid social structure, but lower movement frequency in the higher-density population [51]. Badger populations in Woodchester and Cornwall have historically exhibited higher densities compared to those in Northern Ireland [20, 47, 48]. Hence differences in density could be driving the variation in temporal diffusion metrics observed here, as badgers in less dense populations may range over larger territories searching for resources or mates, leading to higher diffusion values. However, culling is also a critical factor that likely influences badger movement behaviour. Studies have shown that culling disrupts spatial organisation and can significantly increase ranging behaviour, potentially amplifying diffusion metrics [18, 29]. For the Woodchester years included in this study, no density estimates have yet been published but it is likely that culling on neighbouring land reduced badger abundance over time. Similarly, site C4 experienced culling from September 2016 onward, while F1, F2, and C2 had no culling during the study period, and Northern Ireland was subject to prolonged low intensity culling under the TVR strategy. These varying approaches to culling likely contributed to the observed diffusion patterns, highlighting the need to evaluate both population density and impacts of management interventions when interpreting badger movement dynamics.

These findings from the diffusion analysis underscore how short-term movement behaviours can reflect underlying drivers, such as population density and culling. However, while fast-scale metrics like diffusion highlight immediate spatial responses, they do not capture broader patterns in space use or shifts in territorial organisation over time. To fully understand the mechanisms driving the observed movement changes, particularly in response to long-term disturbances like sustained culling, it is essential to also examine the slower temporal dynamics. Analysing long-term patterns provides insight into how badger groups reorganise spatially, adapt to environmental pressures, or experience breakdowns in territorial stability. By integrating both fast and slow timescales, we can better interpret how external pressures reshape movement behaviour not just in the moment, but over months or years.

To explore these longer-term behavioural responses in more detail, the second part of the study focused on slow temporal dynamics, aiming to characterise how badger movement patterns evolve over extended periods. In this context, ‘long-term’ refers to patterns observed over several months, as analysed using EDMD to capture and characterise these dynamics. The main advantage of using EDMD to generate these clusters is that it makes use of the temporal dimension of the data, which considers that a badger can leave its territory and then return again to the same territory. Our results identify spatial regions where metastable clusters remain consistent over time at some study sites. For instance, in the Woodchester data, some clusters exhibit long-term stability throughout the study period. However, in other areas, such as north-east of Woodchester park, substantial changes in the estimated areas were observed, with clusters increasing in size over the period 2018 to 2021 from 0.29 km^2^ to 1.96 km^2^ then decreasing in 2022 to 0.49 km^2^. The merging and splitting of badger group home ranges may suggest a breakdown in territorial behaviour, with increased overlap in the areas used by individuals which formerly had exclusive territories. These changes reflect movement patterns previously observed in populations subjected to culling [18]. We observed a general decrease in the number of clusters for Woodchester (Table 4), accompanied by an increase in cluster area, suggesting that individual animals were ranging over larger areas. This pattern of increasing roaming behaviour and appears to have intensified since the onset of culling in 2018. The observed variations in metastable clusters in Woodchester may therefore be linked to the behavioural responses to the culling around the study area. These findings highlight the potential value of metastable states as a tool for tracking changes in movement dynamics, particularly in response to external interventions such as culling. Unlike simpler methods, such as bait marking or basic measures of ranging behaviour, EDMD allows for the identification of subtle, higher-order patterns in movement dynamics and the transitions between behavioural states over time. This capability provides a more profound understanding of how interventions influence movement behaviours at a system-wide level.

Furthermore, it was observed that the obtained metastable states often overlapped multiple known group ranges, suggesting that badgers were using larger areas than advertised by territorial marking. In the Northern Ireland data set, for example, the average difference in the number of social groups and EDMD clusters over the four years was 3.5, with fewer EDMD clusters on average, and a gradual increase in the number of clusters and group home ranges.

Considering that social groups typically range from 25 to 33 and EDMD clusters from 21 to 28, this represents a moderate change. This difference in spatial organisation may be due to the removal of TB-positive badgers after the first year of the study, which could have changed the social structure and encouraged movement among the remaining badgers. On the other hand, Cornwall sites C2 and C4 had an average difference of −0.3 and −0.75, respectively, showing there was very little difference between the conventional field methods and this approach. For both sites, the number of social groups ranged from 1 to 4, while EDMD clusters ranged from 1 to 4 for C2 and 1 to 6 for C4. These narrow ranges underscore that the differences are minor, reflecting the close agreement between the two approaches. In a few cases, we observed a close match between the number of social groups identified through traditional field methods and the clusters detected by EDMD. This alignment was particularly evident when a single social group dominated the area, as in Cornwall site C4 during the second capture year, where both methods identified only one group. In Cornwall site C2 during the third capture year, only one badger was collared, as work at that site ceased after that year, and while this clearly prevents us from making any broader inferences, we include it to illustrate that EDMD appropriately reflects the minimal movement data available. Meaning that the method detects a single movement trajectory, which is consistent with expectations. Although such instances offer limited insights into group dynamics, they demonstrate the method’s capacity to handle edge cases and confirm that the algorithm does not artificially over-cluster when data are sparse. This occasional alignment between EDMD and field observations suggests that while EDMD has the potential to offer an alternative approach to understanding badger social groups and space use, it may also be capturing different aspects of badger movement dynamics compared to traditional methods. Additionally, discrepancies between the methods could reflect external influences, such as the disruptions caused by culling, which complicate the detection of stable patterns. These findings highlight the complementary nature of the methods and the need to carefully interpret results within the context of external pressures.

Several methodologies have been developed to identify latent or metastable behavioural states from animal movement data, particularly using GPS tracking. Hidden Markov Models (HMMs) are widely used to infer discrete behavioural states based on movement metrics such as step length and turning angle, and they have been applied across diverse taxa [52, 53]. State-space models (SSMs) are another class of models that account for observation errors in tracking data and can infer unobserved behavioural states. They are particularly useful when dealing with noisy data or irregular sampling intervals [54]. Beyond traditional statistical models, unsupervised machine learning techniques have been employed to detect behavioural states without predefined categories. These methods can uncover patterns in movement data that may not be immediately apparent. More recently, unsupervised machine learning approaches such as clustering, change point detection, and autoencoder-based models have been explored for identifying shifts in movement patterns without predefined state assumptions [55]. While these methods typically model behaviour as a sequence of discrete transitions, EDMD offers a complementary perspective by capturing the continuous dynamics and spectral structure underlying movement trajectories. This spectral view allows for the identification of metastable states and transitions without relying on discrete state assumptions, potentially uncovering more nuanced or overlapping behavioural regimes. Situating EDMD within this growing suite of tools highlights its novel contribution to the analysis of animal movement behaviour.

A limitation of our approach stems from the uneven temporal distribution of GPS data, particularly the scarcity of fixes during the first half of the year due to restrictions on trapping during the breeding season to avoid disturbing reproductive females and their offspring [43]. Additionally, the varying lifespan of GPS device batteries contributed to inconsistent data collection, with those in Cornwall averaging 110 days of recording and those in Northern Ireland lasting up to 273 days [19, 20]. Since many collars were deployed in July and August, most devices ceased functioning by the following May, resulting in a lack of data for June and a limited temporal window of data collection. This inconsistency primarily affected our diffusion analysis. Although the generalised linear mixed model accounts for some of this variation through random effects, it cannot fully compensate for periods with sparse or missing observations.

Furthermore, the non-uniform timing of GPS fixes introduced additional inconsistency, impacting our EDMD analysis, which requires equal time intervals between locational fixes. To address this inconsistency, we used linear interpolation. Cornwall’s data required minimal manipulation, as locations were recorded every 20 minutes, whereas Woodchester’s 35-minute intervals and Northern Ireland’s hour intervals necessitated interpolation. While increasing the frequency of fixes could enhance data accuracy, it would also reduce battery life. Additionally, we recognise that a significant number of data points were removed due to the low number of satellites detected (of the overall 19.0% removed for Woodchester, 55.6% of that data was due to the low number of satellites), which could be attributed to the high proportion of woodland (which can obscure GPS satellite connections) in the study areas. This raises the possibility of introducing a systematic bias, as these data exclusions may have disproportionately affected locations nearer the centre of each badger’s range, potentially closer to their main sett. This bias could skew our understanding of badger spatial behaviour, particularly in relation to their core areas of activity. However, Woodroffe et al. [19] (supplementary material) showed that woodland did not affect collar precision in Cornwall, where the same qualitative results were produced when no filters were applied to the data. Despite these limitations and the need to manipulate the data to use some methods, the diffusion results offer a broader understanding of the regular movements of badgers. Furthermore, the EDMD analysis of metastable clusters offers novel insights into long-term badger movement patterns, adding value to the findings even with these constraints on the dataset.

In summary, this study demonstrates the value of combining complementary analytical approaches to understand badger behaviour, particularly their movement and territoriality, across multiple temporal scales. The diffusion analysis captured fine-scale movement dynamics, revealing distinct short-term spatial behaviours that varied across sites and appeared to reflect differences in population density and management interventions such as culling. In parallel, EDMD enabled the identification of spatial clusters corresponding to badger group ranges, offering insights into long-term behavioural organisation and how it may break down under sustained external pressures. In both Northern Ireland and Woodchester Park, discrepancies between EDMD-derived clusters and known social group boundaries likely reflect disruption caused by animal removal, underscoring how population management can reshape territorial structure. At the same time, mismatches with field-derived groupings highlight the need to evaluate the assumptions and limitations of both the analytical methods and the field data.

Together, these analyses present a novel, mathematically grounded framework for interpreting badger spatial behaviour, bridging the gap between short-term movement responses and long-term ecological organisation. The further development and validation of such methods, particularly in the context of wildlife monitoring and management, could be a promising direction for future research. This might include addressing limitations in GPS tracking data (e.g., irregular fix intervals), and exploring the use of Kalman filtering in combination with EDMD to enable more robust system reconstruction from non-uniform time series [56–58].

## Conclusion

This study aimed to investigate badger movement and spatial organisation using GPS tracking data, applying mathematical tools to complement traditional ecological methods. While badger territories have long been studied through approaches such as bait marking [9, 59], the metastability of movement behaviours within populations has remained largely unexplored. By combining diffusion metrics with EDMD, we provide a novel perspective on badger spatial behaviour that captures both fine-scale movement dynamics and broader, longer-term social structures. Diffusion analysis highlights the variability and responsiveness of movement at short timescales, whereas EDMD offers a useful method that captures the social structures that underlie these movements. Together, these methods offer a more comprehensive understanding of territoriality and its disruption under management interventions such as culling.

## Supporting information

**S1 Fig. Maps showing removed data points of badgers isolated from the remaining data (in red) and retained data points for Woodchester (top) and Northern Ireland (bottom)**. Each dot represents a specific date-time location for an individual badger.

**S2 Fig. Mean diffusion (***c***) values (and standard deviation) for male (blue) and female (red) badgers during the study period**. Individual diffusion values are shown by month and sex (circle for male and cross for female). Note, that the low estimation for the diffusion in Woodchester October 2022 comes from 12 data points originating from a single female badger. See Figs 1 and S3 Fig for report of number of fixes and count of badgers per month, respectively.

**S3 Fig. Count of male (blue) and female (red) badgers for each site, per month and year**.

**S4 Fig. Residual plot for the final GLMM model**.

**S5 Fig. The calculated metastable clusters with areas of clusters for A: Northern Ireland, B: Cornwall site C2, C: Cornwall site C4, D: Cornwall site F1, E: Cornwall site F2**. The purple lines are the convex hull of the metastable clusters. Due to the size of the Northern Ireland dataset, the areas of the clusters are not shown. Additionally, for Cornwall sites C2 and C4, there is only one social group present in 2015, so only the convex hull of the EDMD cluster is presented.

**S1 Table. Model selection and comparison**.

**S2 Table. Fixed effect estimates from the GLMM model L** ~ **1+M+Sx+(1** |**Site:Animal)**.

**S1 Appendix. Breakdown of cleaned data**. Reasons why data was removed from the study across sites.

**S2 Appendix. Further statistical analysis**. Univariate statistical analysis of the diffusion data, looking for statistical differences within site, between sex, month and years.

**S3 Appendix. Choice of clusters for EDMD**. Investigation into locating the spectral gap using successive differences and hierarchical clustering.

## Acknowledgments

GL and JRF were partly co-funded by the European Union’s Horizon Europe Project 101136346 EUPAHW. The funders had no role in study design, data collection and analysis, decision to publish, or preparation of the manuscript. JRF was supported by core funding from the University of Surrey for her PhD, and since March 2025, she has been supported by the European Union’s Horizon Europe Programme (Grant No. 101136346, EUPAHW).

For the purpose of Open Access, the author has applied a Creative Commons Attribution (CC BY) public copyright licence to any Author Accepted Manuscript version arising from this submission.

## Data Statement

The code used to analyse the data is available at the Github repository: https://github.com/JFurber/Analysis of Badger Movement.git. The code includes all scripts necessary to reproduce the analysis and results presented in the paper. The analysis was performed using a mixture of R, Python and MATLAB, with specific packages or libraries, which are documented in the repository.

The GPS tracking data of badger movements, collected from Cornwall, are lodged on Movebank (www.movebank.org; Movebank Project 158275131). For the GPS tracking data of badger movements, collected from Woodchester, please contact data access at APHA on. The GPS tracking data of badger movements, collected from Northern Ireland, are lodged on Movebank (www.movebank.org; Movebank Project 5019193945).

