## Supplementary material for "Data-driven analysis of fine-scale badger movement in the UK": S1_Appendix.pdf

### Breakdown of cleaned data

Fig. S6 provides a breakdown of data removal reasons across the study sites. The seven main categories identified are: (i) badger in trap—data entries recorded while the badger was still in the trap (these had already been removed from the Woodchester and Northern Ireland datasets prior to data acquisition); (ii) different capture time—seen in the Northern Ireland dataset, where data from multiple studies with varying timelines were combined; (iii) distance—instances where the Euclidean distance travelled exceeded 26.2 meters per minute multiplied by the time interval between fixes; (iv) DOP—entries with a dilution of precision (DOP) greater than four; (v) duplicates—identical location fixes; (vi) outside of area—locations falling outside the main entries of data (see S1 Fig.); and (vii) satellites—entries recorded with fewer than four satellites. An eighth category, other, includes miscellaneous removal reasons, such as location points recorded in bodies of water.

As noted in the discussion, the primary reason for data removal in Woodchester was a low number of satellites, likely due to the site's dense woodland, which can interfere with GPS signal acquisition and degrade location accuracy (as reflected in higher DOP values). These data points were excluded to maintain consistency with the Cornwall dataset, which had already been pre-processed using similar filtering criteria (see [1]; supplementary materials).

In the Cornwall dataset, which arrived pre-cleaned, the most common reason for data exclusion was excessive movement speed. Specifically, any location where a badger's calculated movement speed exceeded 26.2 meters per minute, the upper bound reported in prior studies [2], was removed, where it was deemed biologically implausible.

For Northern Ireland, the majority of removed data, over 60%, consisted of duplicate entries, likely due to repeated GPS fixes or data merging from multiple sources. This pattern reflects the unique challenges associated with integrating telemetry datasets from different sources.

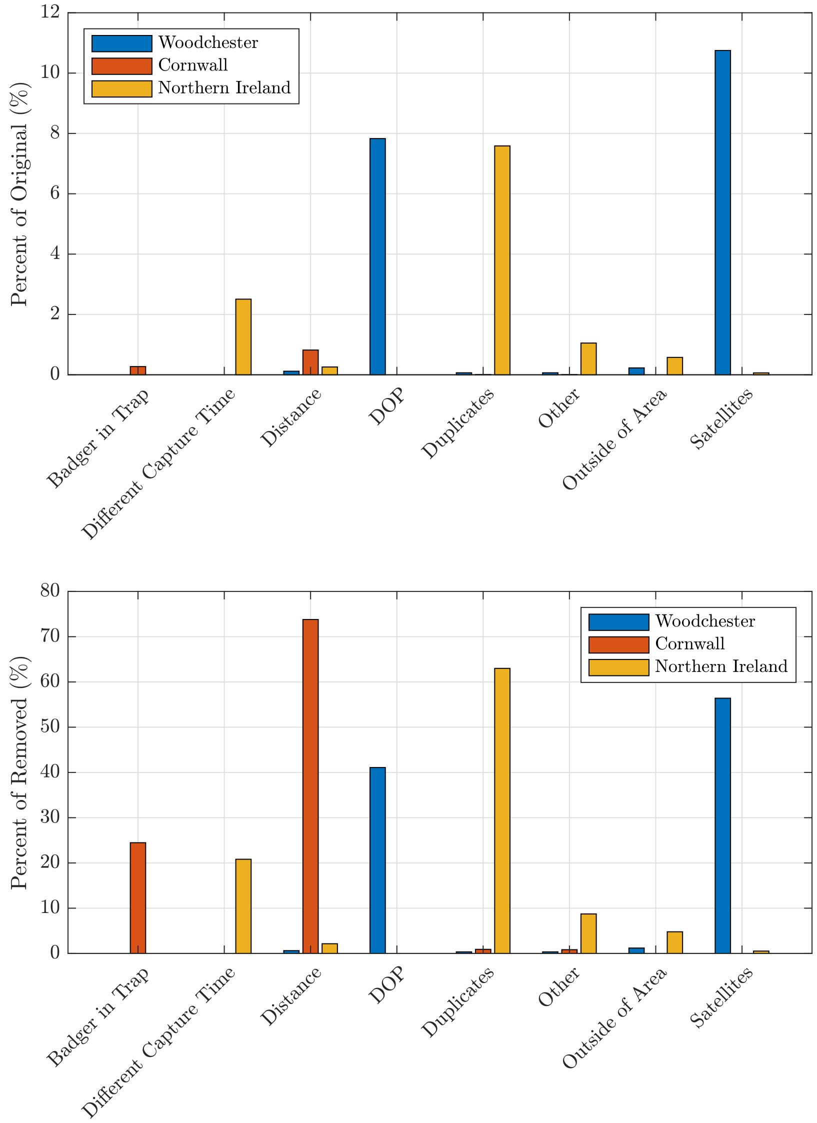


**Fig S6. Summary of data removal reasons for Woodchester, Cornwall (all sites), and Northern Ireland.** The top panel shows each category as a percentage of the total received data, while the bottom panel shows the same categories as a percentage of the removed data.
