## Supplementary material for "Data-driven analysis of fine-scale badger movement in the UK": S1_Table.pdf

**Table S1. Model selection and comparison.**

<table>
<tr><th>Model</th><th>$R^2$ (Adj.)</th><th>AIC</th><th>BIC</th><th>Marginal $R^2$</th><th>Conditional $R^2$</th></tr>
<tr><td>L∼1+M+Sx+(1+ CY|Site)+(1| Site:Animal)*</td><td>0.7910</td><td>80.819</td><td>238.66</td><td>0.1332</td><td>0.7565</td></tr>
<tr><td>L∼1+Sx+(1+M|Site)+(1+ CY|Site)+(1| Site:Animal)</td><td>0.7983</td><td>201.01</td><td>711.35</td><td>0.0262</td><td>0.7667</td></tr>
<tr><td>L∼1+CY+Sx+(1+M|Site)+(1| Site:Animal)</td><td>0.7983</td><td>194.19</td><td>646.56</td><td>0.0338</td><td>0.7591</td></tr>
<tr><td>L∼1+CY+M+Sx+(1| Site:Animal)</td><td>0.7868</td><td>143.22</td><td>243.18</td><td>0.2084</td><td>0.7630</td></tr>
<tr><td>L∼1+CY+M+Sx+(1| Animal)†</td><td>0.7868</td><td>143.22</td><td>243.18</td><td>0.2084</td><td>0.7630</td></tr>
<tr><td>L∼1+CY+M+Sx+(1| Site)‡</td><td>0.4305</td><td>861.59</td><td>961.55</td><td>0.1416</td><td>0.4335</td></tr>
<tr><td>L∼1+CY+M+Sx+(1|Site)+(1| Site:Animal)◕</td><td>0.4305</td><td>863.59</td><td>968.81</td><td>0.1416</td><td>0.4335</td></tr>
<tr><td>L∼1+CY+M+(1| Site:Animal)</td><td>0.7871</td><td>164.24</td><td>258.96</td><td>0.1552</td><td>0.7656</td></tr>
<tr><td>L∼1+M+Sx+(1| Site:Animal)**</td><td>0.7843</td><td>152.82</td><td>231.74</td><td>0.1950</td><td>0.7627</td></tr>
</table>

Capture Year (CY); Month (M); Sex (Sx).

* Final Model, modelling site-specific trends across years and capturing the animal-within-site intercepts.

** Second Model, removing reference to capture year.

†Models each animal's intercepts, but if every animal lives at only one site, that intercept also absorbs all the site-level differences.

‡Models site differences, but assumes all animals at a given site are identical (no extra animal-level scatter).

◕Models variation between sites and the variation between animals within each site.
