## Supplementary material for "Data-driven analysis of fine-scale badger movement in the UK": S2_Appendix.pdf

### Further statistical analysis

We aim to test whether the diffusion values ($c$) differ in each site based on factors such as sex, time (month, year), as well as comparing sites. The first step in analysing the data is to check if the diffusion values follow a normal distribution. We do this using the Shapiro–Wilk test for normality [1], which suggests that the values of $c$ are not normally distributed (W = 0.94446, $p < .001$).

Knowing that the data is not normally distributed, we turn to non-parametric tests that do not assume normality. The first test we apply is the Mann–Whitney U test [2], which is used to compare male and female diffusion values. This test helps us determine whether there is a significant difference between the two groups. In simpler terms, we are asking whether male diffusion differs from female diffusion based on the values of $c$.

Next, we use the Kruskal–Wallis test [3] to compare diffusion values across more than two groups. In this case, we want to examine whether diffusion values differ by months, years, or sites. This test checks if at least one of the groups is significantly different from the others. If the Kruskal-Wallis test reveals a significant difference, we follow up with the Dwass–Steel–Critchlow–Fligner [4] test, a pairwise comparison method. This test helps identify which specific pairs of months, years, or sites have significant differences, providing more detailed insights into where the differences lie. These tests are all computed in R (version 4.5.0) [5] using libraries *dplyr* and *PMCMRplus*, with findings summarised in Table S3.

For Woodchester, under the hypothesis that male diffusion value differs from the female diffusion value, a statistically significant difference between the sexes is observed. Additionally, significant differences are found between years and months, with multiple pairwise comparisons yielding significant results.

Next, for the Cornwall sites, significant sex-based differences in diffusion values are observed only at site C4, while the other three sites do not show significant differences. Analysis of differences between years reveals significant results for sites C2, C4, and F1. When comparing months, significant differences are found at site F2 (though without pairwise significance) and at site F1 (with multiple significant pairwise comparisons).

For the Northern Irish dataset, significant differences in diffusion values are observed between sexes, but no significant differences are found across years or months.

Finally, a Kruskal–Wallis test is performed to compare diffusion values across all sites, yielding a significant result ($\chi^2(5) = 336,\ p < .001$). Pairwise comparisons indicate that diffusion values at Woodchester and all Cornwall sites significantly differ from Northern Ireland ($p < .001$ for C2, C4, F1, F2, and Woodchester). Furthermore, the diffusion values at site C2 were significantly different from C4 ($p = .002$), F2 ($p = .022$), and Woodchester ($p < .001$).

**Table S3. Summary of statistical analysis ($p$-value) for the diffusion of badgers.**

| Site | C2 | C4 | F2 |
|---|---|---|---|
| Sex | $W = 214$ (.905) | $W = 115$ (< .001)$^\star$ | $W = 1052$ (.077) |
| Years | $\chi^2(2) = 10.0$ (.007)$^\star$ | $\chi^2(3) = 11.3$ (.010)$^\star$ | $\chi^2(2) = 5.09$ (.079) |
| *Pairwise Comparisons* | | | |
| Year 1 | vs. Year 3 (.005) | vs. Year 4 (.003) | |
| Months | $\chi^2(9) = 6.83$ (.654) | $\chi^2(9) = 12.5$ (.189) | $\chi^2(11) = 22.6$ (.020)$^\star$ |

| Site | F1 | Woodchester | Northern Ireland |
|---|---|---|---|
| Sex | $W = 2179$ (.401) | $W = 9477$ (.002)$^\star$ | $W = 50139$ (< .001)$^\star$ |
| Years | $\chi^2(4) = 34.6$ (< .001)$^\star$ | $\chi^2(4) = 44.3$ (< .001)$^\star$ | $\chi^2(3) = 1.36$ (.716) |
| *Pairwise Comparisons* | | | |
| Year 1 | vs. Year 3 (.030)<br>vs. Year 4 (.005) | vs. Year 3 (.044)<br>vs. Year 5 (.005) | |
| Year 2 | vs. Year 3 (.023)<br>vs. Year 4 (< .001) | vs. Year 3 (< .001) | |
| Year 3 | vs. Year 4 (.001)<br>vs. Year 5 (.002) | vs. Year 4 (.007)<br>vs. Year 5 (< .001) | |
| Year 4 | vs. Year 5 (< .001) | vs. Year 5 (.006) | |
| Months | $\chi^2(11) = 64.0$ (< .001)$^\star$ | $\chi^2(11) = 43.6$ (< .001)$^\star$ | $\chi^2(11) = 178$ (< .001)$^\star$ |
| *Pairwise Comparisons* | | | |
| Jan | vs. Oct (.011)<br>vs. Nov (.002) | vs. Nov (.043) | vs. Oct (.002)<br>vs. Nov (< .001)<br>vs. Dec (.004) |
| Feb | vs. Aug (.007)<br>vs. Sept (.003)<br>vs. Oct (.002)<br>vs. Nov (.001)<br>vs. Dec (.046) | | vs. Sept (< .001)<br>vs. Oct (< .001)<br>vs. Nov (< .001)<br>vs. Dec (< .001) |
| Mar | | | vs. Sept (.002)<br>vs. Oct (< .001)<br>vs. Nov (< .001)<br>vs. Dec (.004) |
| Apr | | | vs. Oct (< .001)<br>vs. Nov (< .001)<br>vs. Dec (.020) |
| May | vs. Oct (.031)<br>vs. Nov (.016) | | |
| Jun | vs. Oct (.023)<br>vs. Nov (.007) | | |
| Jul | vs. Oct (.037)<br>vs. Nov (.010) | | vs. Sept (.006)<br>vs. Oct (< .001)<br>vs. Nov (< .001)<br>vs. Dec (.009) |
| Aug | vs. Nov (.047) | vs. Oct (.015)<br>vs. Nov (.004) | vs. Sept (.002)<br>vs. Oct (< .001)<br>vs. Nov (< .001)<br>vs. Dec (.009) |
| Sep | | | vs. Oct (< .001)<br>vs. Nov (< .001) |
| Nov | | | vs. Dec (< .002) |

$\star$ $p < .05$, where only significant pairwise comparisons ($p < .05$) are displayed.
