## Supplementary material for "Data-driven analysis of fine-scale badger movement in the UK": S2_Table.pdf

**Table S2. Fixed effect estimates from the GLMM model L∼1+M+Sx+(1| Site:Animal).**

| Predictor | Estimate | Standard Error | t-Statistic | 95% Confidence Interval | $p$-Value |
|---|---|---|---|---|---|
| **Intercept** | 3.5866 | 0.032 | 111.3 | [3.52, 3.65] | $< 0.001$ |
| **Month (Ref: January)** | | | | | |
| February | 0.105 | 0.024 | 4.41 | [0.060, 0.15] | $<0.001$ |
| March | 0.048 | 0.026 | 1.81 | [-0.0041, 0.10] | 0.071 |
| April | 0.033 | 0.030 | 1.09 | [-0.026, 0.092] | 0.23 |
| May | -0.008 | 0.033 | -0.23 | [-0.073, 0.058] | 0.82 |
| June | -0.119 | 0.039 | -3.06 | [-0.19,-0.042] | 0.0023 |
| July | -0.073 | 0.027 | -2.65 | [-0.13, -0.019] | 0.008 |
| August | -0.118 | 0.021 | -5.49 | [-0.16, -0.076] | $<0.001$ |
| September | -0.240 | 0.021 | -11.32 | [-0.28,-0.20] | $<0.001$ |
| October | -0.368 | 0.023 | -16.33 | [-0.41, -0.33] | $<0.001$ |
| November | -0.442 | 0.024 | -18.40 | [-0.49, -0.40] | $<0.001$ |
| December | -0.291 | 0.024 | -12.05 | [-0.34, -0.24] | $<0.001$ |
| **Sex (Ref: Male)** | | | | | |
| Female | -0.201 | 0.041 | -4.86 | [-0.28, -0.12 ] | $<0.001$ |
