## Supplementary material for "Data-driven analysis of fine-scale badger movement in the UK": S3_Appendix.pdf

### Choice of clusters for EDMD

Occasionally, it is not clear where the spectral gap in the spectrum lies. This does not necessarily exclude the usage of spectral methods for identifying the metastable states of a system. So, for an order of $N$ eigenvalues, $\lambda_1 > \lambda_2 > \ldots > \lambda_N$, then a spectral gap is defined as the first $\lambda_i$ such that Eq. (6) is satisfied.

To support the identification of this gap, we attempt two different approaches. We first plot the eigenvalues and their successive differences. The first local maximum in difference, followed by a significant drop, is taken as a preliminary indicator of the spectral gap.

The second approach involves constructing a dendrogram (middle panel) using hierarchical clustering (Ward's method) on the eigenvalues, treating each as a one-dimensional data point. In essence, hierarchical clustering combines similar values step by step, based on how close they are to each other. In this case, similarity is measured using Euclidean distance between eigenvalues. A fixed threshold (0.01) is then applied horizontally to the dendrogram to identify significant gaps between eigenvalues. This grouping offers a complementary heuristic to the eigenvalue difference plot for identifying the spectral gap and estimating the number of metastable states.

The resulting groups of eigenvalues are reflected in the colour-coded scatter plot (right panel), where a solid line marks the spectral gap determined from the difference plot (left panel). In most cases, the dendrogram-based and difference-based methods agree. For example, in site C2, the dendrogram suggests four clusters, while the difference method highlights seven key eigenvalues. This process is inherently heuristic, and while there is no universally accepted method for selecting the number of significant eigenvalues, our combined approach provides a consistent and interpretable framework across datasets.

**Table S4. Number of clusters selected using two spectral gap methods for each site and year.**

| | Total | | Year 1 | | Year 2 | | Year 3 | | Year 4 | | Year 5 | |
|---|---|---|---|---|---|---|---|---|---|---|---|---|
| | De | Di | De | Di | De | Di | De | Di | De | Di | De | Di |
| Woodchester | 8 | 8 | 11 | 12 | 9 | 9 | 11 | 11 | 5 | 5 | 10 | 7 |
| Northern Ireland | 11 | 11 | 14 | 21 | 21 | 21 | 14 | 28 | 12 | 25 | | |
| C2 | 4 | 7 | 3 | 4 | 2 | 4 | 1 | 1 | | | | |
| C4 | 6 | 6 | 5 | 6 | 1 | 1 | 4 | 4 | 2 | 2 | | |
| F1 | 5 | 5 | 5 | 5 | 5 | 5 | 4 | 4 | 2 | 2 | 1 | 1 |
| F2 | 8 | 8 | 6 | 7 | 5 | 9 | 3 | 5 | | | | |

Dendrogram-based (De) with a maximum difference of 0.01 between clusters within the dendrogram.
Based on the first largest difference (Di) between consecutive eigenvalues.
Figure S7 illustrates these methods on the full dataset for each site.

### Woodchester

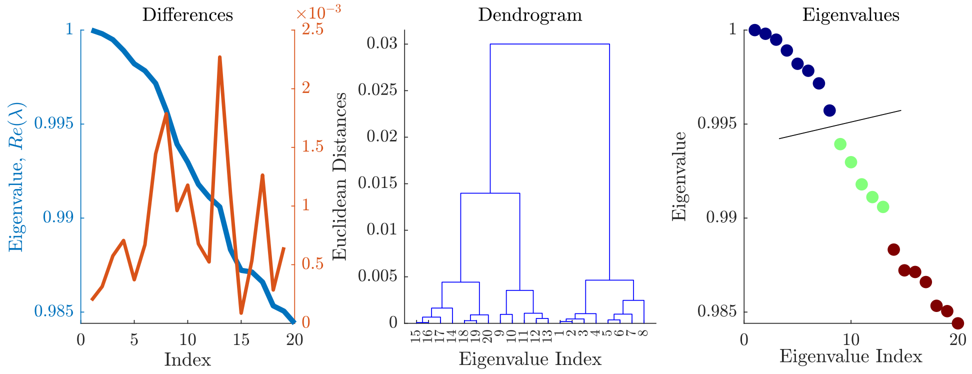


### Northern Ireland

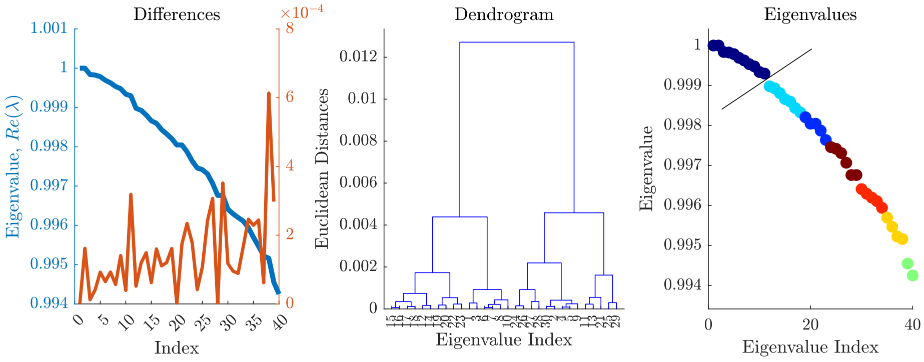


## C2

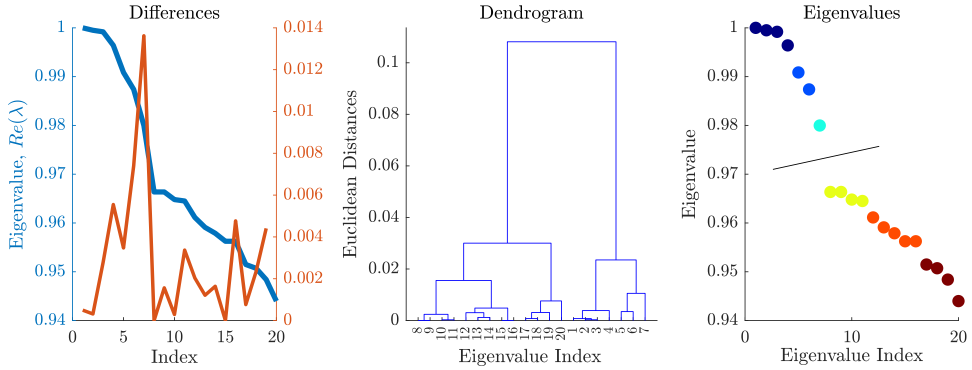

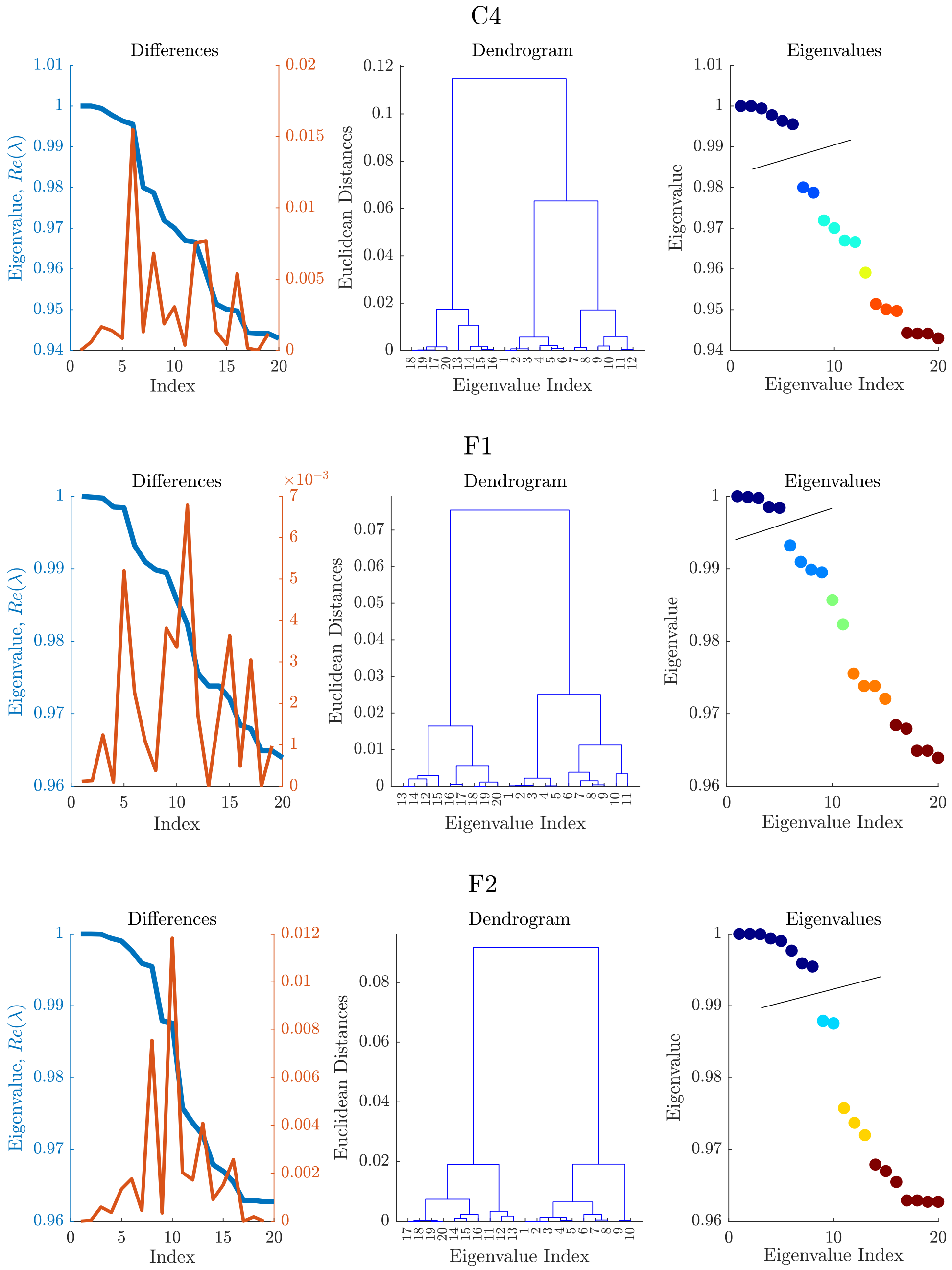


**Fig S7.** Comparing the methods to choose the spectral gap based on the differences in the eigenvalues and through the use of a dendrogram. The result of the optimal choice seen in the third figure, with the line relating to the calculated differences and the colours relating to the dendrogram.
