## Supplementary figures and images for "Data-driven analysis of fine-scale badger movement in the UK"

### S1_Fig.pdf

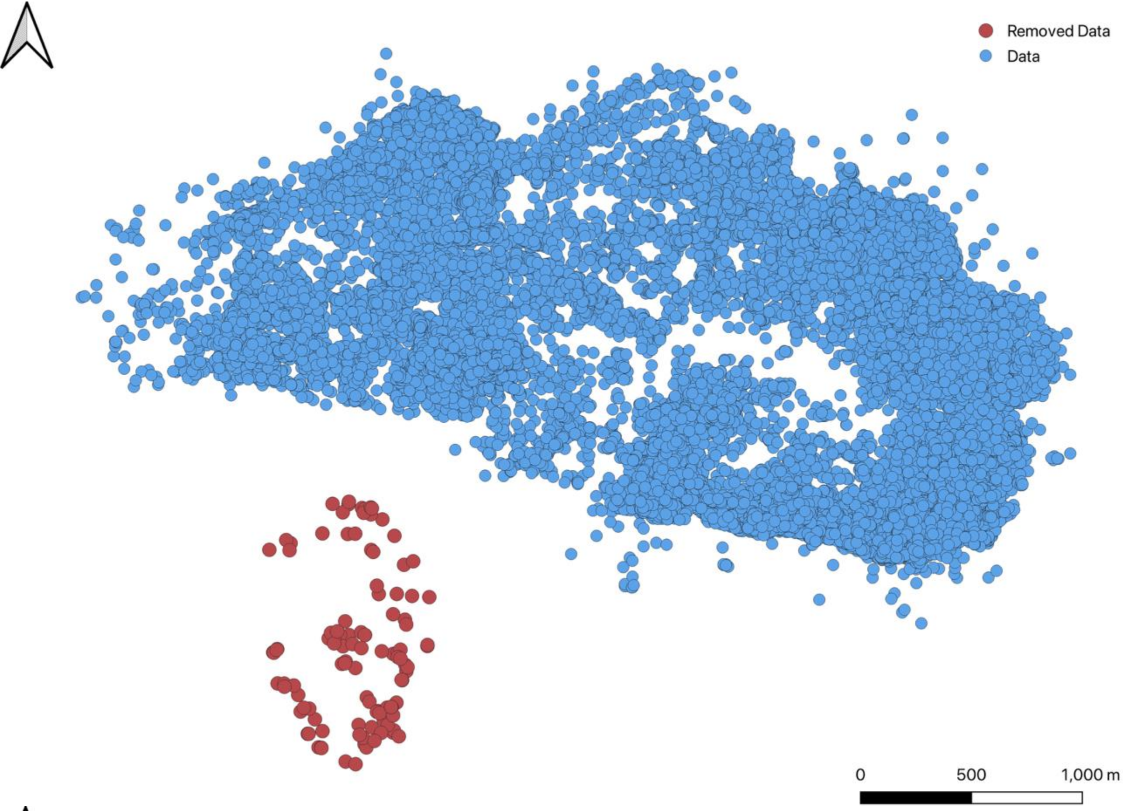

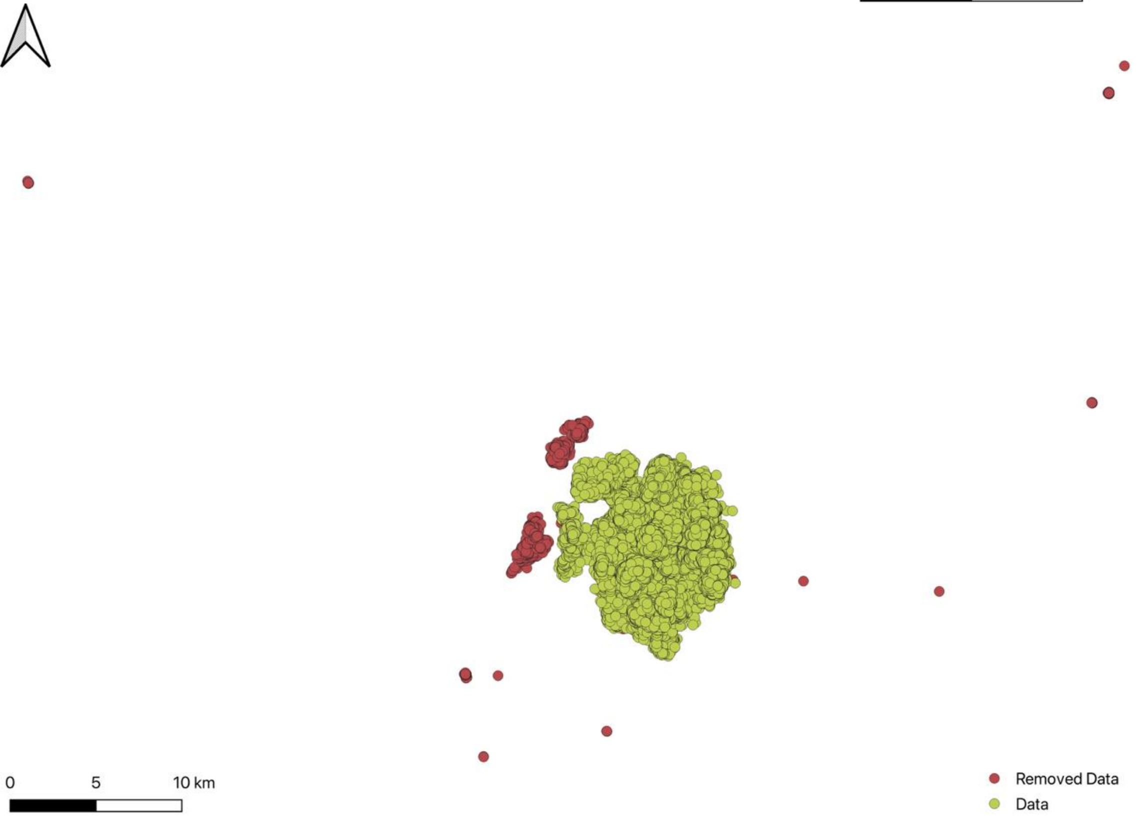

### S2_Fig.pdf

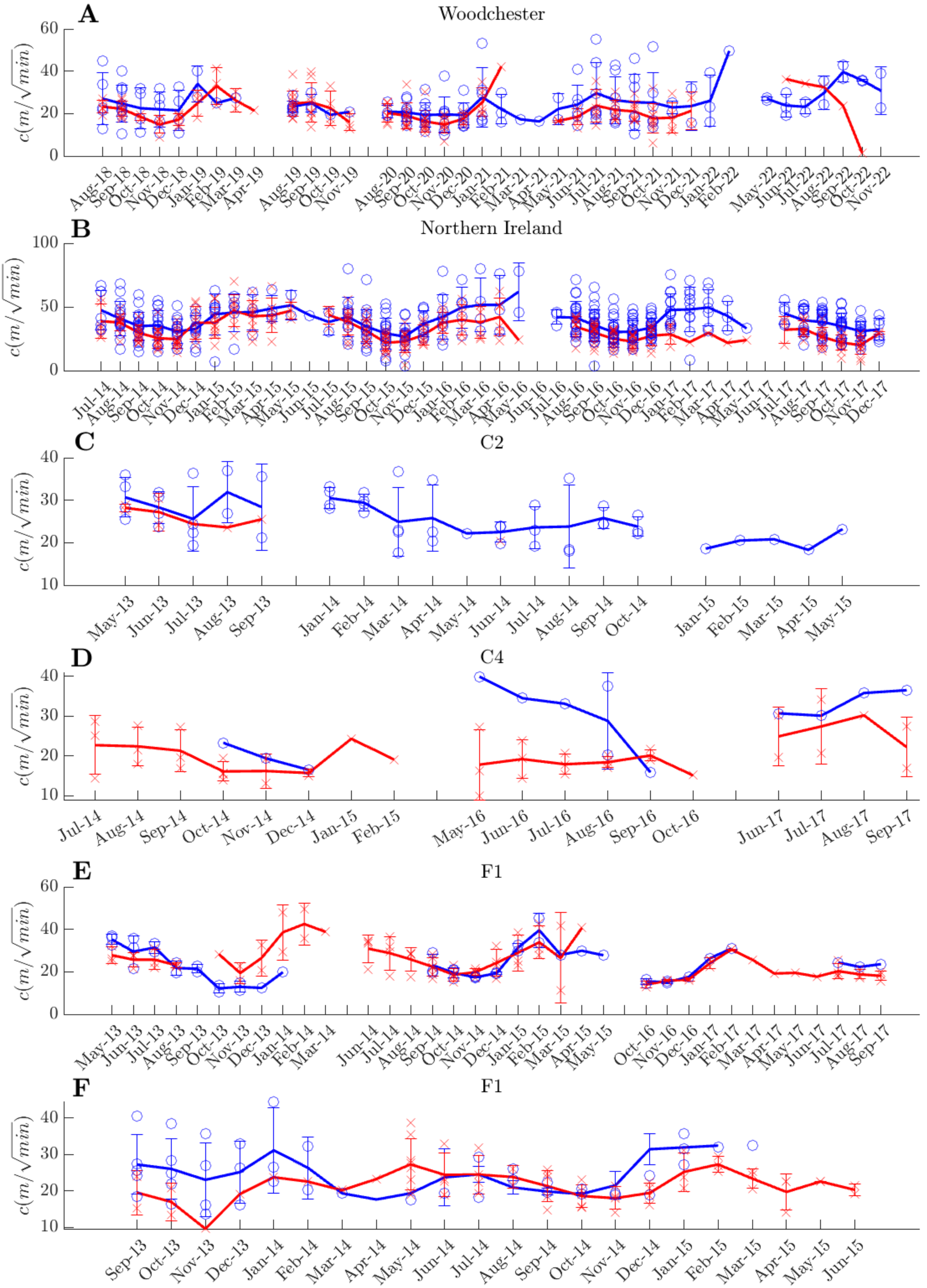

### S3_Fig.pdf

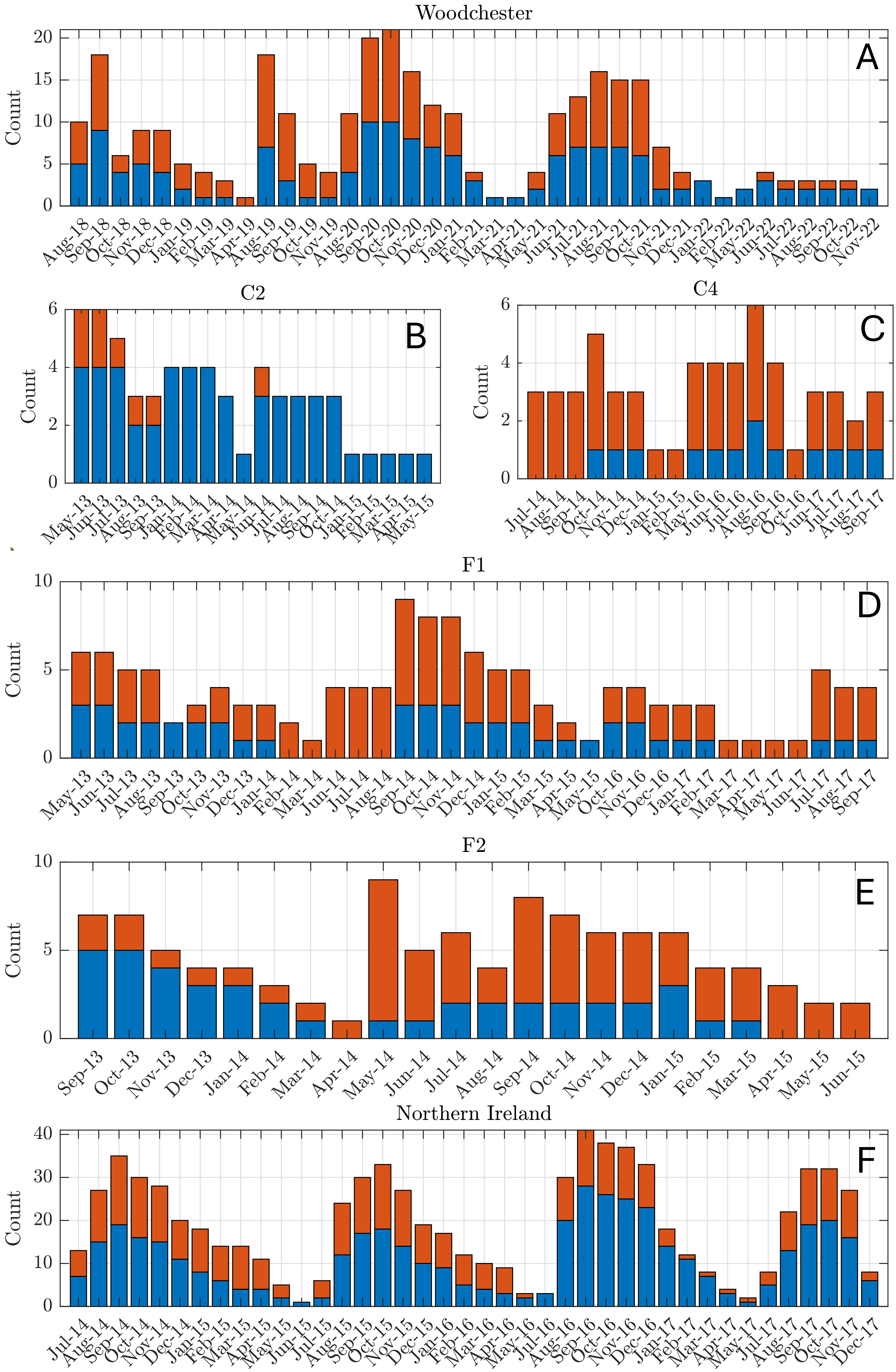

### S4_Fig.pdf

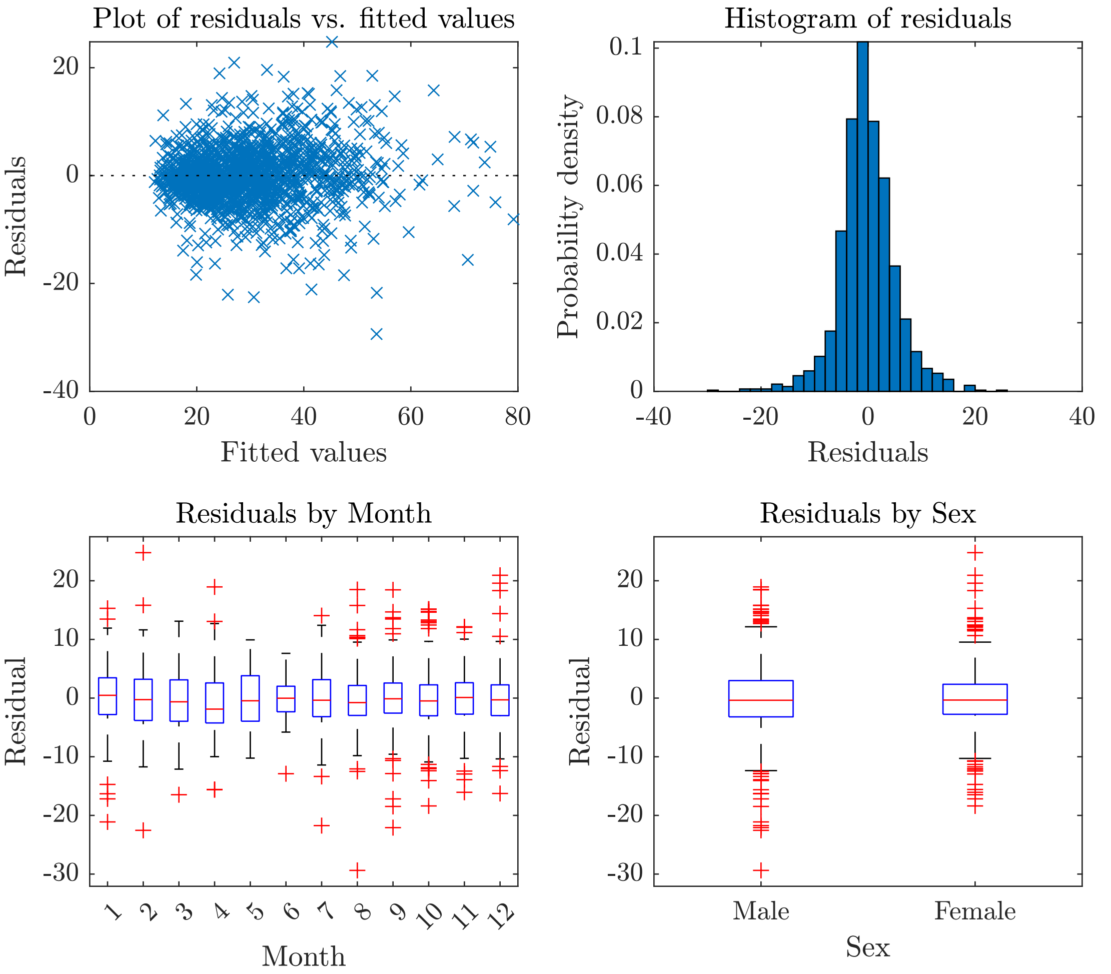

### S5_Fig.pdf

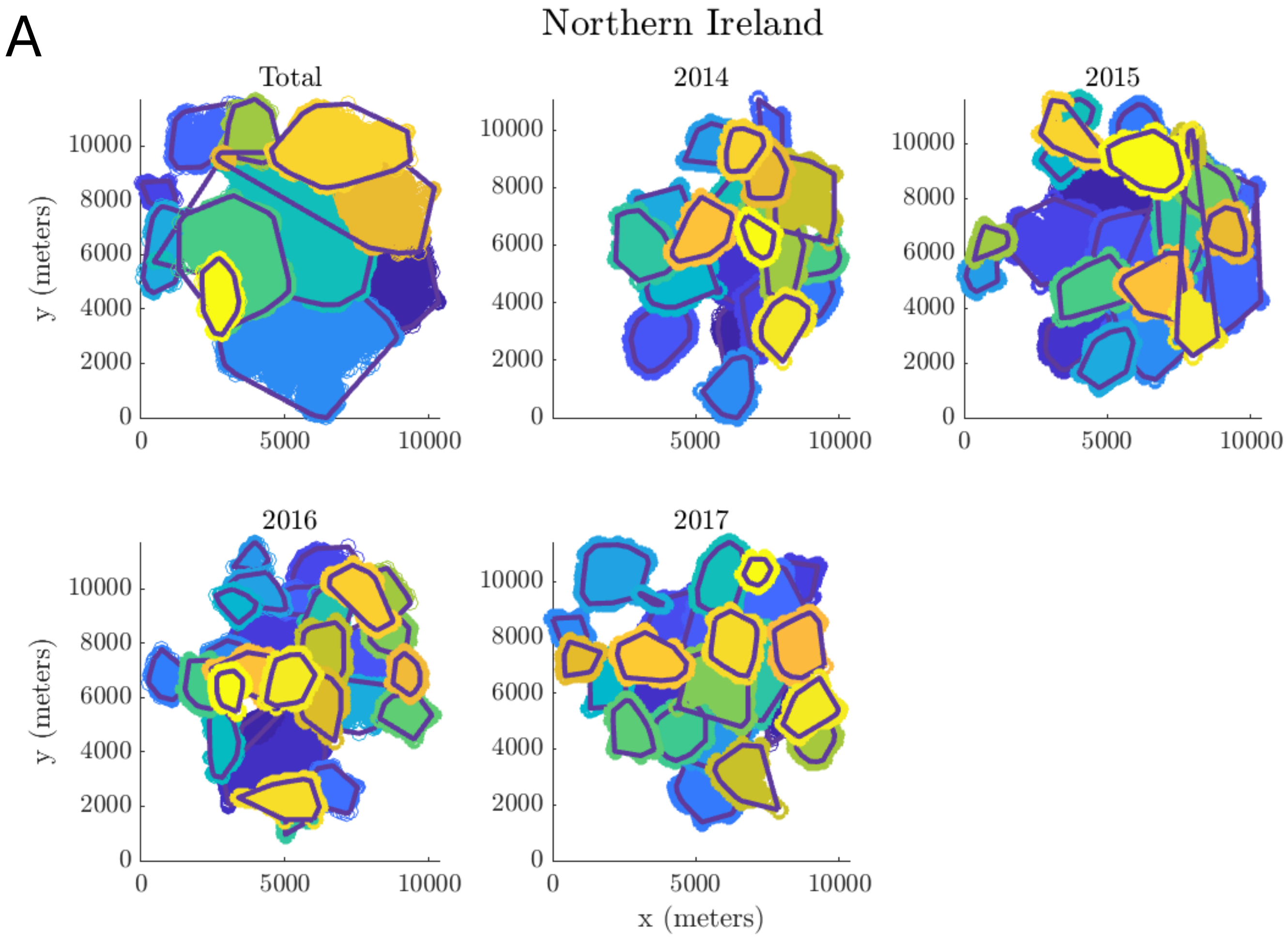

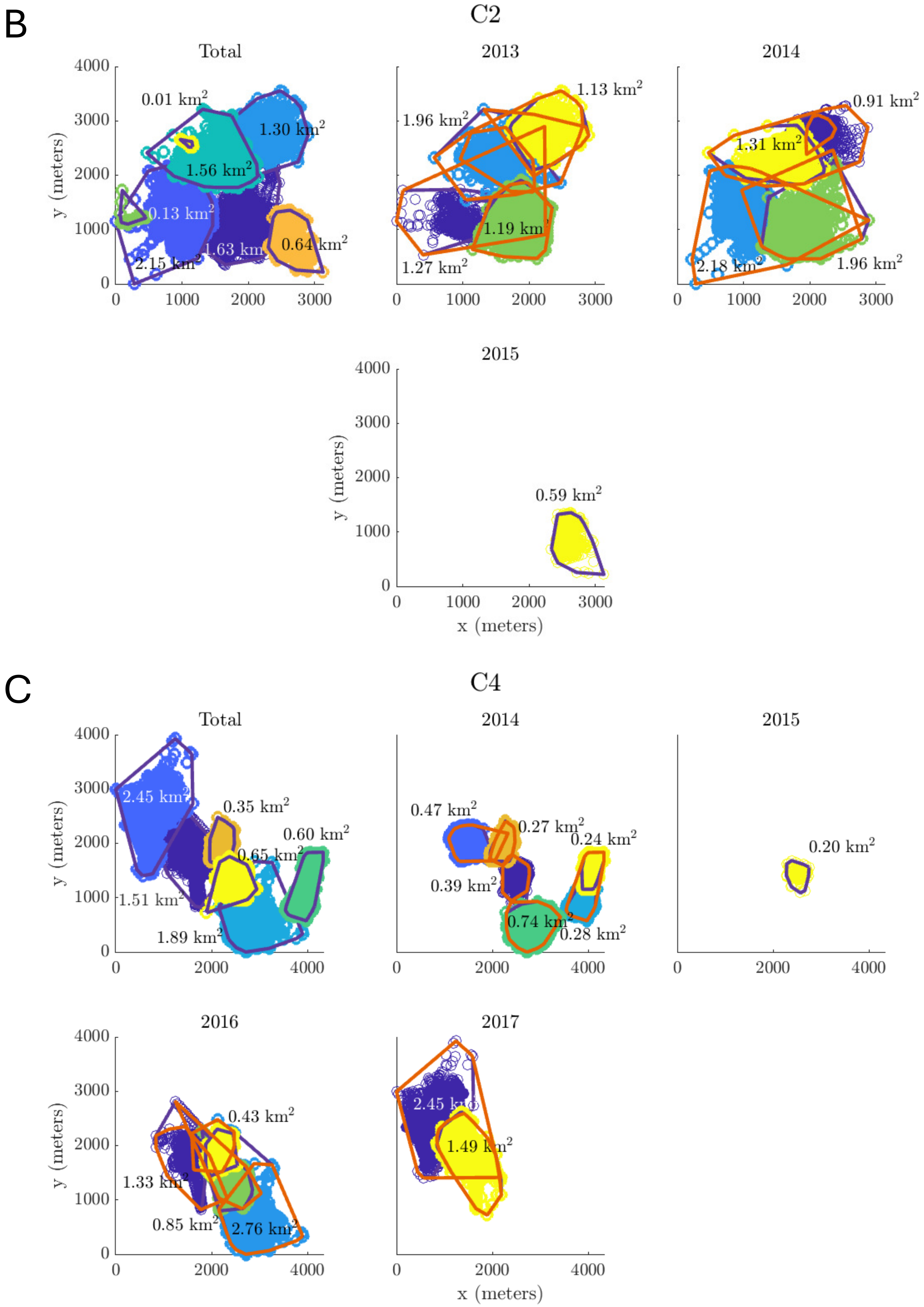

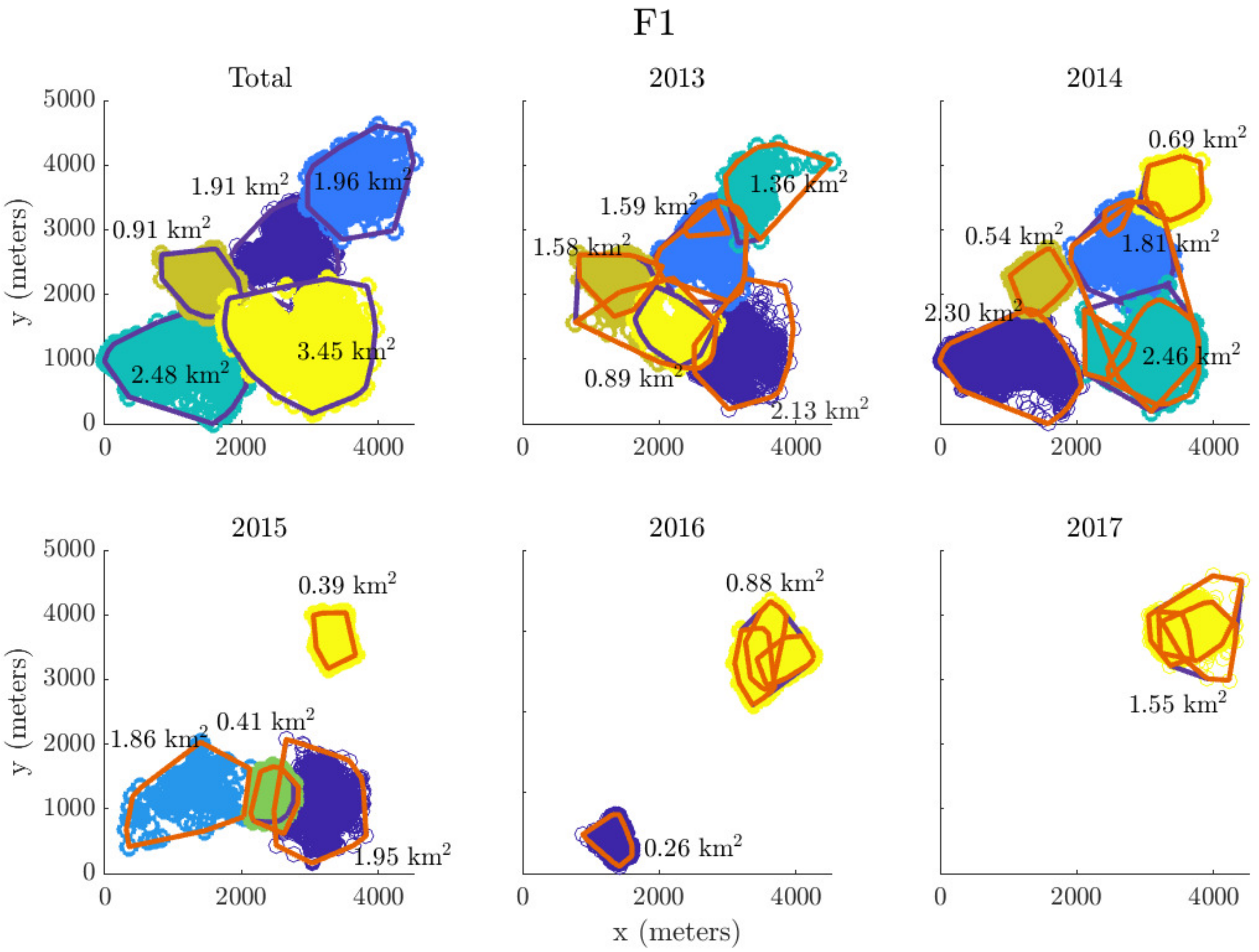

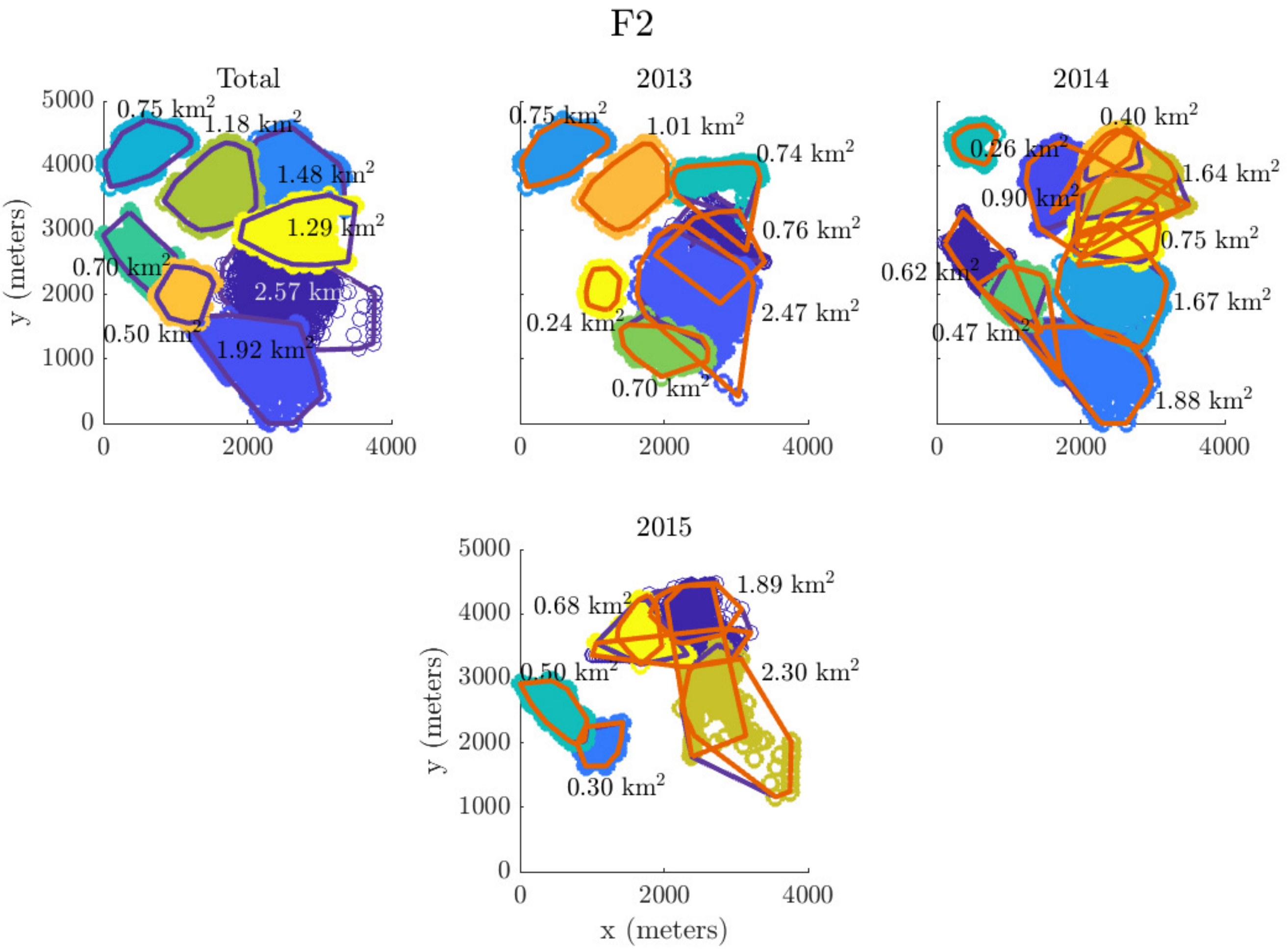
